# Poly(A)-tail-length-dependent surveillance of viral RNA by PABPC1 orchestrates broad-spectrum antiviral defense

**DOI:** 10.64898/2026.01.12.699136

**Authors:** Shimin Yang, Li Zhou, Xiaoya Huang, Shunhua Zhu, Weiyi Yu, Xiao Guo, Jintao Liu, Jikai Deng, Yuzhen Zhang, Jiejie Liu, Qianyun Liu, Ming Guo, Wenjie Xiang, Xin Wang, Zhen Zhang, Zhixiang Huang, Ke Lan, Hongyun Wang, Yu Chen

## Abstract

RNA processing and modification are critical for virus replication and pathogenesis, yet how the host exploits viral RNA features—particularly the poly(A) tail—for antiviral defense remains unclear. Through multi-omics integration and systematic functional screening, we identify the poly(A)-binding protein PABPC1 as a broad-spectrum restriction factor against multiple coronaviruses. We demonstrate that PABPC1 preferentially binds viral RNAs bearing short poly(A) tails—distinct from the longer and more heterogeneous host poly(A) tails—and, in a poly(A)-length–dependent manner, recruits the mitochondrial exonuclease EXD2 to assemble a degradative RNA–protein complex. Both the recognition and subsequent degradation of viral RNA strictly depend on poly(A) tail length, enabling selective decay of short-tailed viral transcripts while sparing host mRNAs. Importantly, inflammatory signaling enhances the expression of PABPC1 and EXD2, suggesting its role as an inducible host defense pathway activated during viral infection. Capitalizing on this discovery, we engineered a synthetic fusion protein mimicking the PABPC1–EXD2 complex and achieved efficient delivery using lipid nanoparticles (LNPs). This rationally designed therapeutic exhibits robust suppression of coronavirus replication in both cellular and murine models. Collectively, our findings uncover a novel host antiviral strategy that targets a conserved viral RNA structural element and provide a conceptual framework for developing host-derived and broad-spectrum anti-coronavirus therapeutics.

## Introduction

Coronaviruses such as SARS-CoV-2, murine hepatitis virus (MHV), and human coronavirus OC43 (HCoV-OC43) exploit host RNA-binding proteins (RBPs) and mRNA regulatory pathways to facilitate viral genome replication and translation^1–3^. These positive-sense RNA viruses carry a single-stranded RNA genome with a 5’ cap and a 3’ polyadenylated tail, features that allow them to mimic host mRNAs and engage host translation machinery efficiently^4^. While viral replication requires hijacking of host RBPs, infected cells simultaneously mount antiviral defenses, including RNA degradation, interferon responses, and cytokine signaling^5–7^. Understanding how host RNA regulatory factors respond to coronavirus infection is therefore essential for elucidating viral pathogenesis and host defense mechanisms.

Despite increasing knowledge of host-virus interactions, the mechanisms connecting coronavirus replication to host RNA turnover remain incompletely understood. Poly(A)-binding protein cytoplasmic 1 (PABPC1) is a central regulator of eukaryotic mRNA stability and translation^8^. PABPC1 comprises four non-identical RNA-recognition motifs (RRMs) at the N-terminus and a C-terminal region containing a proline-rich segment followed by a globular PABC domain^9^. By binding to the polyadenylated tails of mRNAs, PABPC1 protects transcripts from degradation, promotes translation initiation through interactions with the translation machinery, and modulates mRNA decay via recruitment of deadenylases and decay factors^10,11^.

PABPC1 exhibits dual roles across diverse RNA virus infections. Several viruses exploit PABPC1 to enhance their own gene expression: PABPC1 promotes respiratory syncytial virus (RSV) mRNA translation^12^, and facilitates cap-dependent translation of influenza A virus (IAV) mRNAs by binding the viral 5’ UTR, thereby enabling IAV to resist host translational shutoff^13^. PABPC1 also interacts with the nucleocapsid (N) protein of porcine epidemic diarrhea virus (PEDV) to support PEDV replication, and its depletion profoundly impairs viral RNA synthesis and viral titers^14^. In contrast, PABPC1 knockdown increases the stability and accumulation of RNAs from Kaposi’s sarcoma–associated herpesvirus (KSHV)^15^. During infection with bovine coronavirus (BCoV), PABPC1 competes with the viral N protein for binding to the poly(A) tail, thereby suppressing viral mRNA translation^16^. Despite these diverse functional roles, studies dissecting PABPC1 activity in human coronaviruses remain limited.

Poly(A) tails play a critical role in viral RNA replication. In poliovirus, classical studies have established that a minimum poly(A) length of approximately 12 nucleotides is required to support efficient negative-strand RNA synthesis and productive infection^17,18^. Similarly, in coronaviruses, the 3’ poly(A) tail is indispensable for optimal RNA replication, with tail length directly influencing the kinetics and overall efficiency of genome amplification^19^. Viral genomes that lack a poly(A) tail or carry substantially shortened tails exhibit markedly impaired or delayed replication^20^. However, how viral poly(A) tails are selectively recognized by host RNA-binding proteins, and how such recognition dictates the fate of viral RNAs, remains largely unresolved.

EXD2 is a 3’-5’ exonuclease implicated in nuclear and cytoplasmic RNA processing, DNA repair, and RNA quality control^21^. Although both PABPC1 and EXD2 have well-characterized roles in RNA metabolism, whether and how they cooperate to regulate viral RNA decay has not been explored.

Here, we report that PABPC1 and EXD2 functionally cooperate to promote viral RNA degradation. Using SARS-CoV-2 as a model, we show that PABPC1 binds viral RNA in a largely sequence-independent manner, while EXD2, through its 3’-5’ exonuclease domain, facilitates RNA decay. Loss of either factor prolongs viral RNA half-life and enhances replication of SARS-CoV-2, MHV, and HCoV-OC43, but not viruses lacking poly(A) tails such as Zika virus, indicating a poly(A)-dependent mechanism. Both PABPC1 and EXD2 are dynamically upregulated by inflammatory signaling, with transcript and protein levels increasing upon stimulation with IL-1β, TNFα, or LPS in human cells, and in SARS-CoV-2-infected patient samples and K18-hACE2 mouse lungs. Guided by these insights, we engineered a synthetic fusion protein linking PABPC1’s RRM1-4 domains to EXD2’s exonuclease domain, which exhibits potent, dose-dependent inhibition of SARS-CoV-2 replication, outperforming either protein alone. Formulated as mRNA in lipid nanoparticles (LNPs), this therapeutic suppresses viral replication in vitro and reduces viral loads in vivo, highlighting a poly(A)-dependent antiviral pathway that can be exploited for nucleic acid-based interventions.

Together, these findings establish a previously unrecognized RNA-degradation axis, wherein PABPC1 and EXD2 cooperate to restrict coronavirus replication in a poly(A)-dependent manner and are dynamically regulated by inflammatory cues. Our study provides mechanistic insight into host-mediated viral RNA decay and highlights the potential of engineered PABPC1-EXD2 fusion mRNA as a novel antiviral therapeutic platform.

## Results

### PABPC1 is identified as an RNA-binding protein of positive-sense RNA (+ssRNA) viruses through multi-omics analysis and high-throughput screening

To investigate RNA-binding proteins (RBPs) of positive-sense RNA (+ssRNA) viruses, we first analyzed nine independently reported datasets of SARS-CoV-2 RBPs^1,2,22–26^ (**Fig. 1A-B**). To reduce inter-study variability, proteins present in at least four of the nine datasets were selected, yielding 94 high-confidence RBPs (**Fig. 1C**). Among them, PABPC1 appeared in all nine datasets, making it the most consistently identified RBP. We next compared these 94 high-confidence SARS-CoV-2 RBPs with those reported for HCoV-OC43, dengue virus (DENV), and rhinovirus (RV) to determine whether these factors represent broadly utilized RBPs across +ssRNA viruses^2,27^. 69 proteins were shared, representing 10.4% of the total RBPs for these viruses (**Fig. 1D**). Further integration of five genome-wide screens and literature curation on these 69 shared proteins yielded 49 RBPs that significantly affect SARS-CoV-2 replication^28,29^ (**Fig. 1E**). Most function as antiviral host factors, with PABPC1 exhibiting antiviral activity in four datasets. Protein-protein interaction (PPI) network and GO enrichment analyses of the 49 RBPs revealed clustering into mRNA metabolism and cytoplasmic stress granules, and involvement in pathways including stress granule assembly, RNA splicing, RNA metabolism, and translation regulation (**fig. S1**). Notably, PABPC1 participates in multiple pathways, highlighting its key host role. To experimentally validate the bioinformatic predictions, we employed the previously established and widely used SARS-CoV-2 ΔN-GFP-HiBiT replicon delivery particles (RDPs) to assess the effects of these RBPs on viral replication^30–33^ (**Fig. 1F**). Three siRNAs were designed for each of the 49 RBPs, and knockdown efficiency was confirmed by qPCR after transfection into Caco-2-N cells (**fig. S2, table S1**). qPCR measurement of SARS-CoV-2 envelop (E) and Open Reading Frame 1ab(ORF1ab) expression showed that knockdown of most genes enhanced RDPs replication, while PABPC1 demonstrated the most significant role in regulating viral replication (**Fig. 1G-H**).

**Fig. 1.**
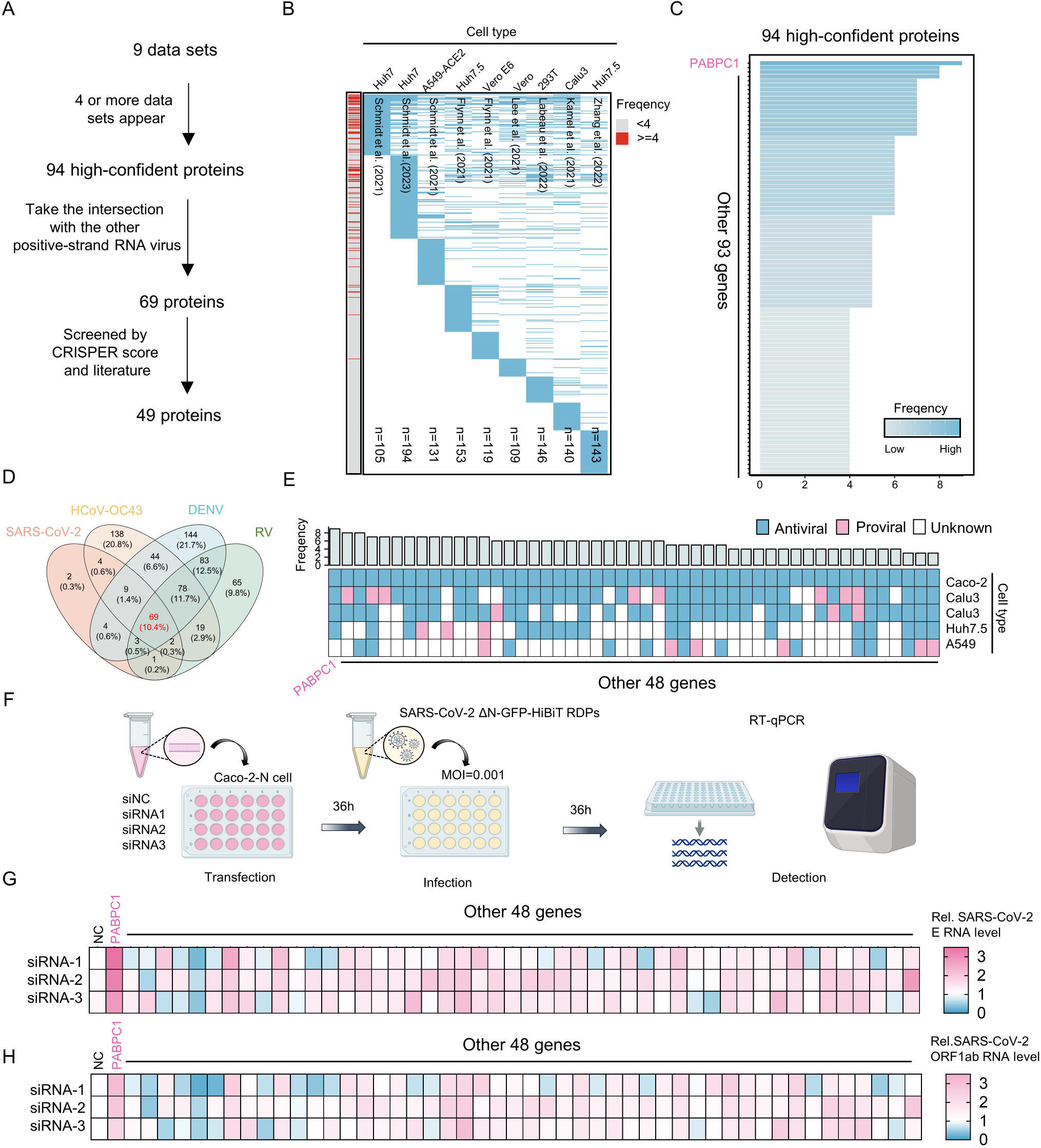
PABPC1 is identified as a RNA-binding protein of positive-sense RNA (+ssRNA) viruses through multi-omics analysis and high-throughput screening (A) Schematic overview of the multi-omics and high-throughput screening strategy used to identify SARS-CoV-2 RNA-binding proteins (RBPs). (B) Heatmap of SARS-CoV-2 RNA-binding proteins across nine independent datasets, proteins detected in four or more datasets are highlighted in red. (C) Bar plot of 94 high-confidence RBPs. (D) Venn diagram showing shared and virus-specific RNA-binding proteins among SARS-CoV-2, HCoV-OC43, DENV, and RV. (E) Heatmap of integration of five genome-wide screens and literature curation identifies 49 RBPs that significantly affect SARS-CoV-2 replication. (F) Schematic of RBPs knockdown and SARS-CoV-2 ΔN-GFP-HiBiT replicon delivery particles (RDPs) infection workflow. (G-H) Knockdown of selected RBPs in Caco-2-N cells followed by qPCR measurement of SARS-CoV-2 E and ORF1ab expression.

### PABPC1 restricts the replication of SARS-CoV-2, HCoV-OC43 and MHV, but not ZIKV

To investigate the role of PABPC1 across +ssRNA viruses, we examined SARS-CoV-2, HCoV-OC43, MHV, and ZIKV. We first validated efficient PABPC1 knockdown across all corresponding cell lines (**fig. S3A-D**). Loss of PABPC1 markedly enhanced SARS-CoV-2 RDPs replication in Caco-2-N cells (**Fig. 2A**), HCoV-OC43 replication in RD cells (**Fig. 2B**), and MHV replication in Neuro2a cells (**Fig. 2C**). In contrast, ZIKV replication was largely unaffected, likely because ZIKV lacks a 3’ poly(A) tail^34^, supporting a poly(A)-dependent antiviral role of PABPC1. To test this hypothesis, we performed RNA immunoprecipitation followed by qPCR (RIP-qPCR) after viral infection. Western blot confirmed efficient immunoprecipitation of PABPC1 (**fig. S3E**). RIP-qPCR analysis revealed that PABPC1 binds viral RNA from SARS-CoV-2, HCoV-OC43, and MHV without apparent sequence preference, whereas ZIKV RNA was not enriched (**Fig. 2E-H**). This likely reflects the absence of a 3’ poly(A) tail in ZIKV, which diminishes PABPC1 binding affinity, consistent with the minimal impact of PABPC1 depletion on ZIKV replication. Collectively, these results indicate that loss of PABPC1 preferentially enhances the replication of +ssRNA viruses possessing a poly(A) tail.

**Fig. 2.**
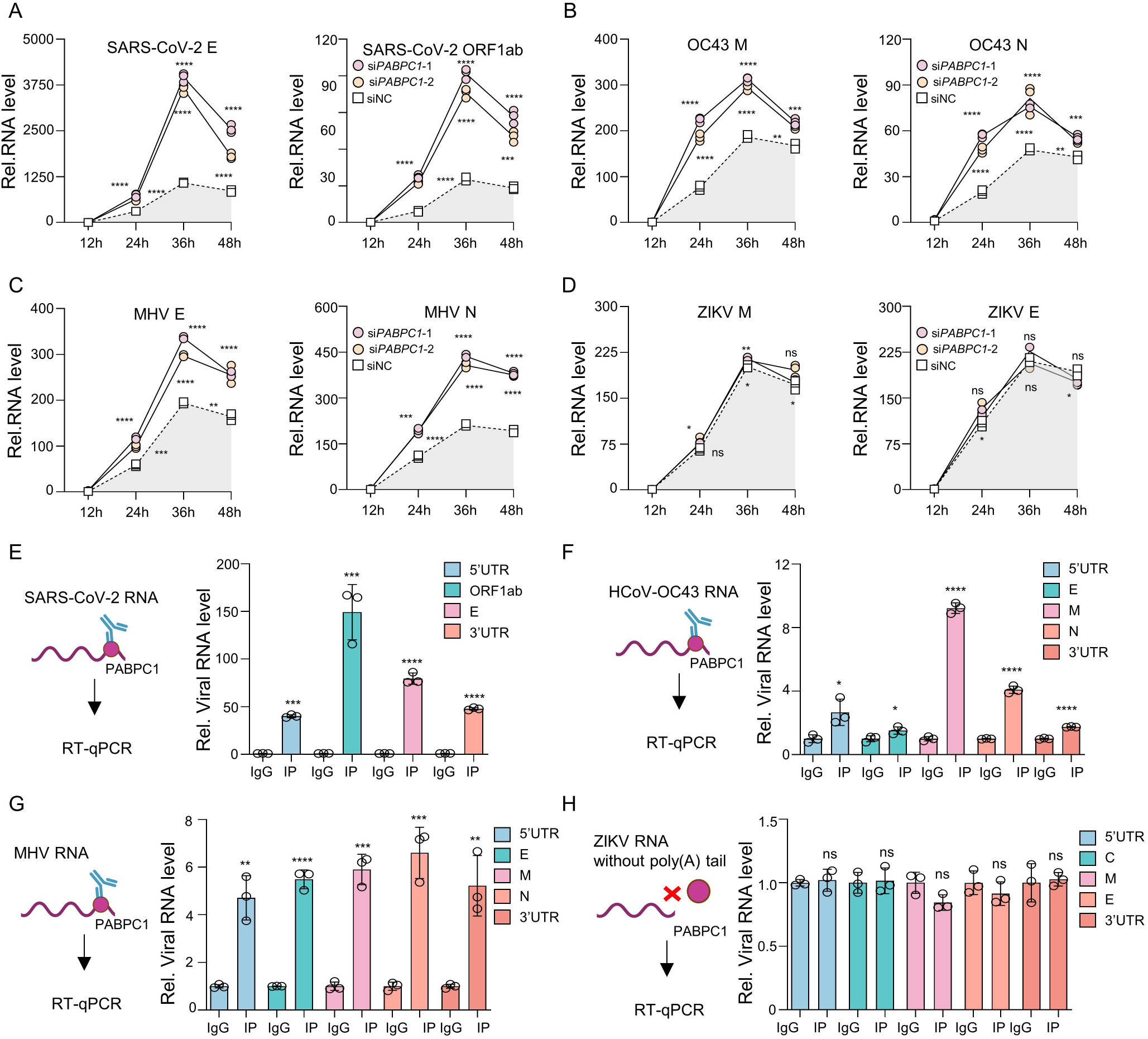
PABPC1 restricts the replication of SARS-CoV-2, HCoV-OC43 and MHV, but not ZIKV. (A) qPCR analysis of SARS-CoV-2 E and ORF1ab RNA level at 12, 24, 36, and 48 h post-infection in Caco-2-N cells with PABPC1 knockdown. (B) qPCR analysis of HCoV-OC43 M and N RNA level at 12, 24, 36, and 48 h post-infection in RD cells with PABPC1 knockdown. (C) qPCR analysis of MHV E and N RNA level at 12, 24, 36, and 48 h post-infection in Neuro2a cells with PABPC1 knockdown. (D) qPCR analysis of ZIKV M and E RNA level at 12, 24, 36, and 48 h post-infection in A549 cells with PABPC1 knockdown. (E) RIP-qPCR analysis of SARS-CoV-2 gene enrichment in IgG and PABPC1 immunoprecipitated samples following SARS-CoV-2 RDPs infection. (F) RIP-qPCR analysis of HCoV-OC43 gene enrichment in IgG and PABPC1 immunoprecipitated samples following HCoV-OC43 infection. (G) RIP-qPCR analysis of MHV gene enrichment in IgG and PABPC1 immunoprecipitated samples following MHV infection. (H) RIP-qPCR analysis of ZIKV gene enrichment in IgG and PABPC1 immunoprecipitated samples following ZIKV infection. Data are representative of three independent experiments and were analyzed by two-tailed unpaired t test. Graphs show the mean ± SD (n = 3) derived from three independent experiments. NS, not significant for *P* > 0.05, *\*P* < 0.05, *\*\*P* < 0.01, *\*\*\*P* < 0.001.

### PABPC1 depletion enhances coronavirus replication by binding to viral poly(A) tails

To investigate the mechanism by which PABPC1 restricts +ssRNA viruses possessing a poly(A) tail, we next focused on SARS-CoV-2, a globally significant pathogen responsible for substantial morbidity, mortality, and socio-economic impact^35,36^. We conducted further tests under the condition of live virus infection of SARS-CoV-2. Consistently, PABPC1 knockdown significantly increased the SARS-CoV-2 N and E RNA levels at 24 h and 48 h post infection (**Fig. 3A-C**). We next generated PABPC1 knockout Caco-2 and Huh7 cell lines using CRISPR/Cas9 (**Fig. 3D-E**) and observed robust elevation of SARS-CoV-2 N and E RNA levels both intracellularly and extracellularly (**Fig. 3F-I**). Similarly, PABPC1 knockout markedly increased SARS-CoV-2 N protein levels (**Fig. 3J-K**). We then performed RNA-seq analysis to determine the global transcriptional impact of PABPC1 knockout. Pearson correlation heatmaps and PCA analysis demonstrated high within-group similarity, indicating strong stability and consistency across biological replicates (**fig. S4A-B**). Transcriptome sequencing further revealed a marked increase in viral read-mapping rates, together with a global upregulation of all viral genes in PABPC1-deficient samples, indicating substantially enhanced SARS-CoV-2 replication (**Fig. 3L-M**). Differential expression analysis (Fold change > 2; *P*< 0.05) identified 932 upregulated and 501 downregulated host genes upon PABPC1 deletion, demonstrating its broad regulatory influence on the host transcriptome (**fig. S4C**). Notably, inflammatory cytokines were predominantly upregulated, suggesting that enhanced SARS-CoV-2 replication in PABPC1-deficient cells may be accompanied by an elevated inflammatory response (**fig. S4D**). GO enrichment of upregulated genes revealed significant enrichment in processes related to immune response, double-stranded RNA binding, and components such as the MLL3/4 complex and P-bodies. Consistently, KEGG analysis showed activation of the TNF, MAPK, and NF-κB signaling pathways (**fig. S4E**). In contrast, downregulated genes were enriched in lipid and oxoacid metabolic processes, peptidase regulator activity, protease binding, and cellular components including the endoplasmic reticulum lumen and blood microparticles. KEGG analysis further indicated suppression of pathways related to complement and coagulation cascades, glycolysis/gluconeogenesis, and tyrosine metabolism (**fig. S4F**). These findings indicate that loss of PABPC1 enhances SARS-CoV-2 replication, thereby amplifying host immune responses and perturbing cellular energy metabolism.

**Fig. 3.**
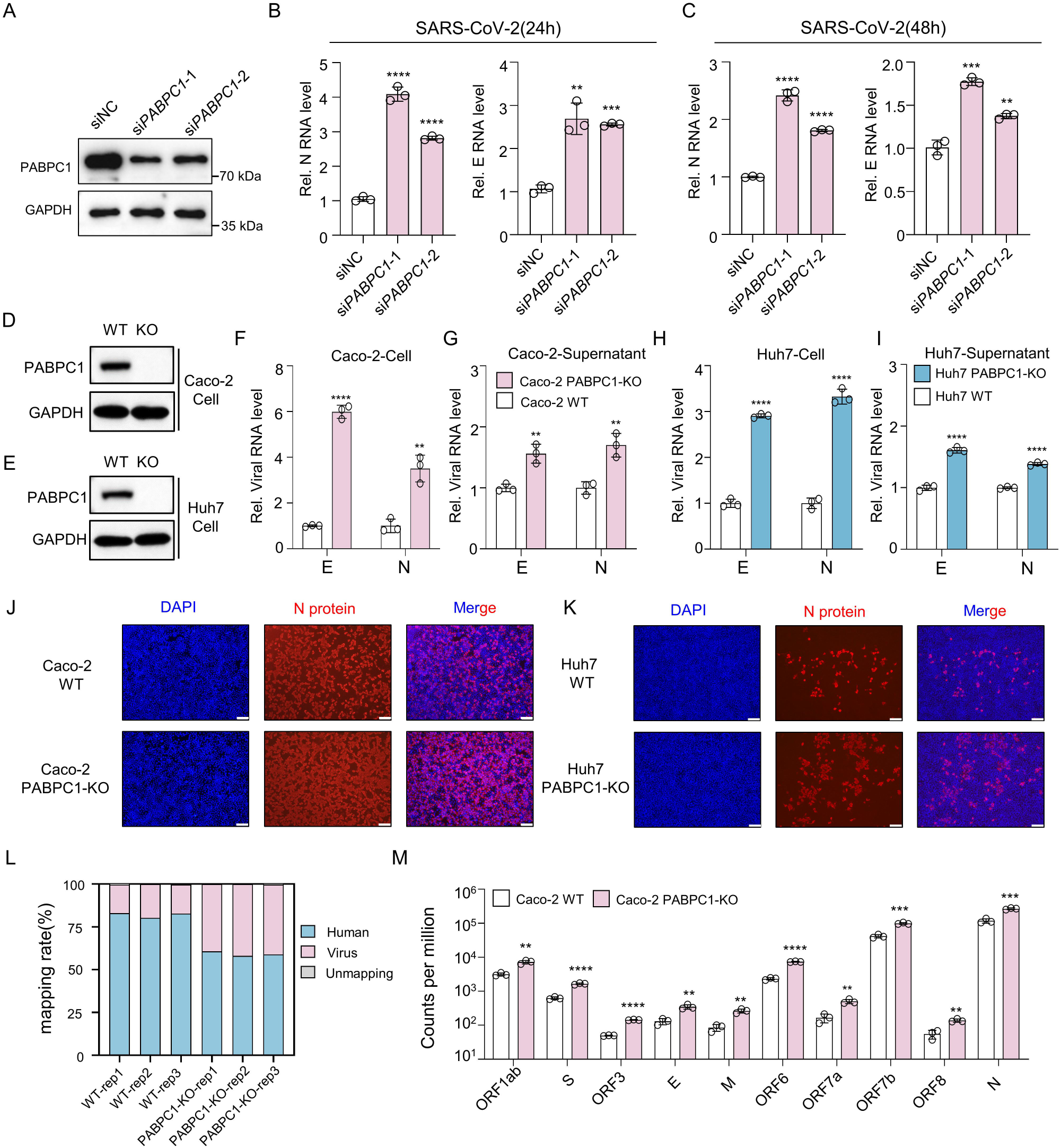
PABPC1 depletion enhances coronavirus replication in vitro. (A) Western blot analysis confirming PABPC1 knockdown in Caco-2 cells. Cells were subsequently infected with SARS-CoV-2 at an MOI = 0.02, and viral RNA levels of the N and E genes were quantified by qPCR at 24 h (B) and 48 h (C) post-infection. Western blot analysis confirming PABPC1 knockout in Caco-2 cells (D) and Huh7 cells (E). (F-I) qPCR analysis of RNA levels of N or E in cell or supernatant from wild-type Caco-2 or PABPC1 knockout cells and wild-type Huh7 or PABPC1 knockout cells, with infection by SARS-CoV-2 at an MOI = 0.02 for 24 h. (J-K) Immunofluorescence analysis of N protein (red) in wild-type Caco-2 or PABPC1 knockout cells and wild-type Huh7 or PABPC1 knockout cells, with infection by SARS-CoV-2. Scale bar, 100 μm. (L) RNA-seq analysis of SARS-CoV-2 and human genome mapping rates in wild-type and PABPC1 knockout Caco-2 cells, with infection by SARS-CoV-2. (M) Bar plot showing counts per million (CPM) of individual SARS-CoV-2 genes in wild-type and PABPC1 knockout Caco-2 cells. Data are representative of three independent experiments and were analyzed by two-tailed unpaired t test. Graphs show the mean ± SD (n = 3) derived from three independent experiments. NS, not significant for *P* > 0.05, \**P* < 0.05, \*\**P* < 0.01, \*\*\**P* < 0.001.

Based on previous reports that 1,10-Phen and ML324 competitively inhibit PABPC1–poly(A) binding^37^, we evaluated their effects on SARS-CoV-2 replication (**Fig. 4A**). CCK-8 assays in Caco-2-N cells showed no detectable cytotoxicity at concentrations below 0.5 μM for 1,10-Phen and 4 μM for ML324 (**fig. S5A-B**). Pre-treatment of Caco-2-N cells with 1,10-Phen or ML324 prior to SARS-CoV-2 RDPs infection significantly increased SARS-CoV-2 E and ORF1ab gene levels in a dose-dependent manner (**fig. S5C-F**). Consistent with the SARS-CoV-2 RDPs results, pretreatment of Caco-2 cells with either inhibitor for 24 h followed by infection (MOI = 0.02) significantly increased SARS-CoV-2 E and N gene levels (**Fig. 4B-C**). Similarly, both inhibitors enhanced infection by the related coronavirus HCoV-OC43 (**fig. S5G-H**), whereas no significant effect was observed on ZIKV replication (**fig. S5I-J**), likely because ZIKV lacks a canonical poly(A) tail, and thus does not rely on PABPC1–poly(A) interactions.

**Fig. 4.**
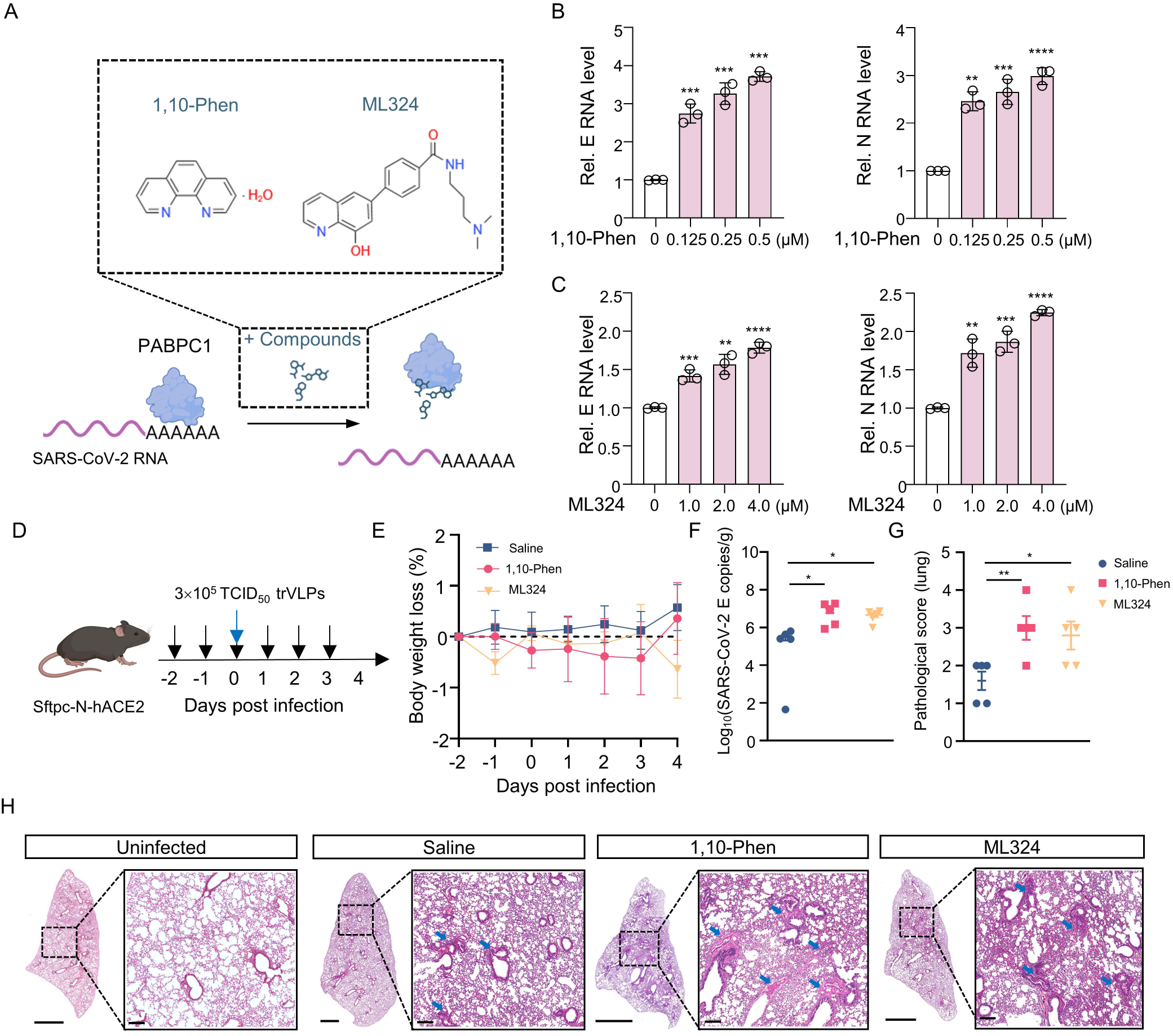
Pharmacological inhibition of PABPC1–poly(A) interaction enhances coronavirus replication in vitro and in vivo. (A) Schematic representation of small-molecule inhibitors 1,10-Phen and ML324 competitively binding to PABPC1. (B-C) qPCR analysis of SARS-CoV-2 E and N RNA levels in Caco-2-N cells pretreated with 1,10-Phen or ML324 for 24 h, followed by infection with SARS-CoV-2 ΔN-GFP-HiBiT RDPs at an MOI of 0.001. (D) Schematic representation of Sftpc-N-hACE2 mice pretreated with 1,10-Phen or ML324, followed by intranasal infection with SARS-CoV-2 RDPs. Black arrows indicate intraperitoneal injection of 1,10-Phen or ML324, and blue arrows indicate intranasal administration of SARS-CoV-2 RDPs. (E) Line graph showing body weight changes in Sftpc-N-hACE2 mice. Mice were treated with saline (control) or the small-molecule inhibitors 1,10-Phen or ML324 prior to SARS-CoV-2 RDPs infection, n=5. (F) qPCR analysis of SARS-CoV-2 E RNA levels in the lung of Sftpc-N-hACE2 mice, n=5. (G) Pathology scores of lungs from saline-treated, 1,10-Phen-treated, and ML324-treated mice, n=5. (H) Representative H&E-stained lung sections from uninfected, saline-treated, 1,10-Phen-treated, and ML324-treated mice, n=5. Scale bar, 2000 and 200 µm. Scale bar, 2000 and 200 µm. Data are representative of three independent experiments and were analyzed by two-tailed unpaired t test. Graphs show the mean ± SD (n = 3 in B-C, n = 5 in E-G) derived from three independent experiments. NS, not significant for *P* > 0.05, *\*P* < 0.05, *\*\*P* < 0.01, *\*\*\*P* < 0.001.

We further evaluated their antiviral activity in vivo using Sftpc-N-hACE2 mice, a transgenic model previously generated in our laboratory that stably expresses SARS-CoV-2 N protein and human angiotensin-converting enzyme 2 (ACE2) in the lung. This model has been applied for studies involving RDPs-mediated SARS-CoV-2 infection^38^. Mice were intraperitoneally injected with 1 mg/kg 1,10-Phen or 2.5 mg/kg ML324 for three consecutive days before intranasal infection with 3 × 10 TCID SARS-CoV-2 RDPs, followed by continued daily administration (**Fig. 4D**). While body weights remained stable (**Fig. 4E**), qPCR analysis of mouse lungs showed that viral E gene copies were markedly elevated in the inhibitor-treated groups (**Fig. 4F**). Lung pathology was evaluated using a standardized scoring system ranging from 0 to 5. A score of 0 indicated normal alveolar architecture; 1 reflected mild inflammatory cell infiltration without alveolar wall thickening; 2 indicated moderate infiltration with slight wall thickening; 3 corresponded to pronounced inflammatory aggregation and regional thickening; and scores of 4-5 represented extensive inflammatory cell accumulation, marked alveolar wall thickening, bronchiolar obstruction, and lung consolidation. Consistent with these criteria, histological analysis revealed markedly exacerbated lung injury in inhibitor-treated mice, characterized by alveolar wall thickening, dense inflammatory infiltration, and exudate accumulation, leading to significantly higher pathology scores compared with saline-treated controls (**Fig. 4G-H**). Taken together, the loss of PABPC1 markedly enhances SARS-CoV-2 replication both *in vitro* and *in vivo*, and this effect is closely associated with its poly(A)-binding activity.

### PABPC1 directly binds coronavirus RNA and selectively accelerates the decay of short poly(A)-tailed viral transcripts

To examine whether its RNA-binding profile changes upon infection, we performed RIP-seq. Pearson correlation and PCA analyses confirmed high reproducibility among replicates (**fig. S6A-B**). Before SARS-CoV-2 RDPs infection, both IP and Input samples showed almost exclusively host-derived reads. Strikingly, after infection, viral read-mapping was markedly enriched in the IP samples, increasing from ∼4.8% to ∼17.6%. These results indicate that SARS-CoV-2 infection drives a pronounced shift in PABPC1 binding toward viral RNA (**Fig. 5A**). We further analyzed the RIP-seq profiles and observed that, in SARS-CoV-2 RDPs-infected cells, PABPC1 IP samples displayed markedly increased peak signals across the entire viral genome (**Fig. 5B**). This indicates that PABPC1 preferentially associates with SARS-CoV-2 RNA upon infection but shows no positional preference along the viral genome. We further confirmed this interaction using an in-vitro EMSA assay, in which purified in-vitro–transcribed viral SARS-CoV-2 RNA was incubated with recombinant prokaryotically expressed PABPC1 protein. The assay revealed clear mobility shifts, demonstrating a direct physical interaction between PABPC1 and viral RNA (**Fig. 5C**). To determine whether PABPC1 affects the stability of SARS-CoV-2 RNA, we transfected a plasmid expressing the viral M gene into Caco-2 WT or PABPC1-KO cells and subsequently treated cells with actinomycin D to block transcription. qPCR analysis showed that loss of PABPC1 markedly increased the half-life of M RNA, indicating that PABPC1 promotes viral RNA decay (**Fig. 5D**). To determine whether this PABPC1-mediated RNA decay is linked to poly(A) tail dynamics, we next performed Tail-seq to assess the impact of PABPC1 on viral RNA poly(A) tail length. Pearson correlation and PCA analyses confirmed high reproducibility among replicates (**fig. S6C-D**). The results showed that loss of PABPC1 led to elongation of SARS-CoV-2 RNA poly(A) tails (**Fig. 5E**). Detailed analysis revealed that host RNAs generally have longer and more heterogeneous poly(A) tails, whereas viral RNAs are concentrated around 50-60 nt (**Fig. 5F**). Using *in vitro*-transcribed RNAs bearing defined poly(A) lengths, we transfected these RNAs into control or PABPC1-depleted HEK293T cells and monitored RNA abundance 12 h post-transfection. Interestingly, the effect of PABPC1 depletion on RNA stability varies depending on the length of the poly(A) tail. Notably, PABPC1 knockdown led to a pronounced increase in the stability of the 54A-tailed RNA. Moreover, as the poly(A) tail length increases, PABPC1 depletion progressively reduces RNA stability (**Fig. 5H**). These findings indicate that PABPC1 directly binds coronavirus RNA and shortens its half-life in a poly(A)-tail-length–dependent manner, with the balance between RNA degradation and stabilization varying according to tail length. To determine the functional domains required for PABPC1-mediated restriction, we next dissected its domain architecture using a SARS-CoV-2 luciferase reporter system. We generated a luciferase construct containing the 5’ and 3’ UTRs of SARS-CoV-2 to mimic viral RNA in cells and enable quantitative detection (**fig. S6E**). Luciferase assays demonstrated that PABPC1 suppresses SARS-CoV-2-Luc activity in a dose-dependent manner (**fig. S6F**). Truncation analysis further revealed that this inhibitory effect is strictly dependent on the RNA-binding modules RRM1–4, indicating that PABPC1’s ability to engage viral RNA is essential for its antiviral function (**fig. S6G**).

**Fig. 5.**
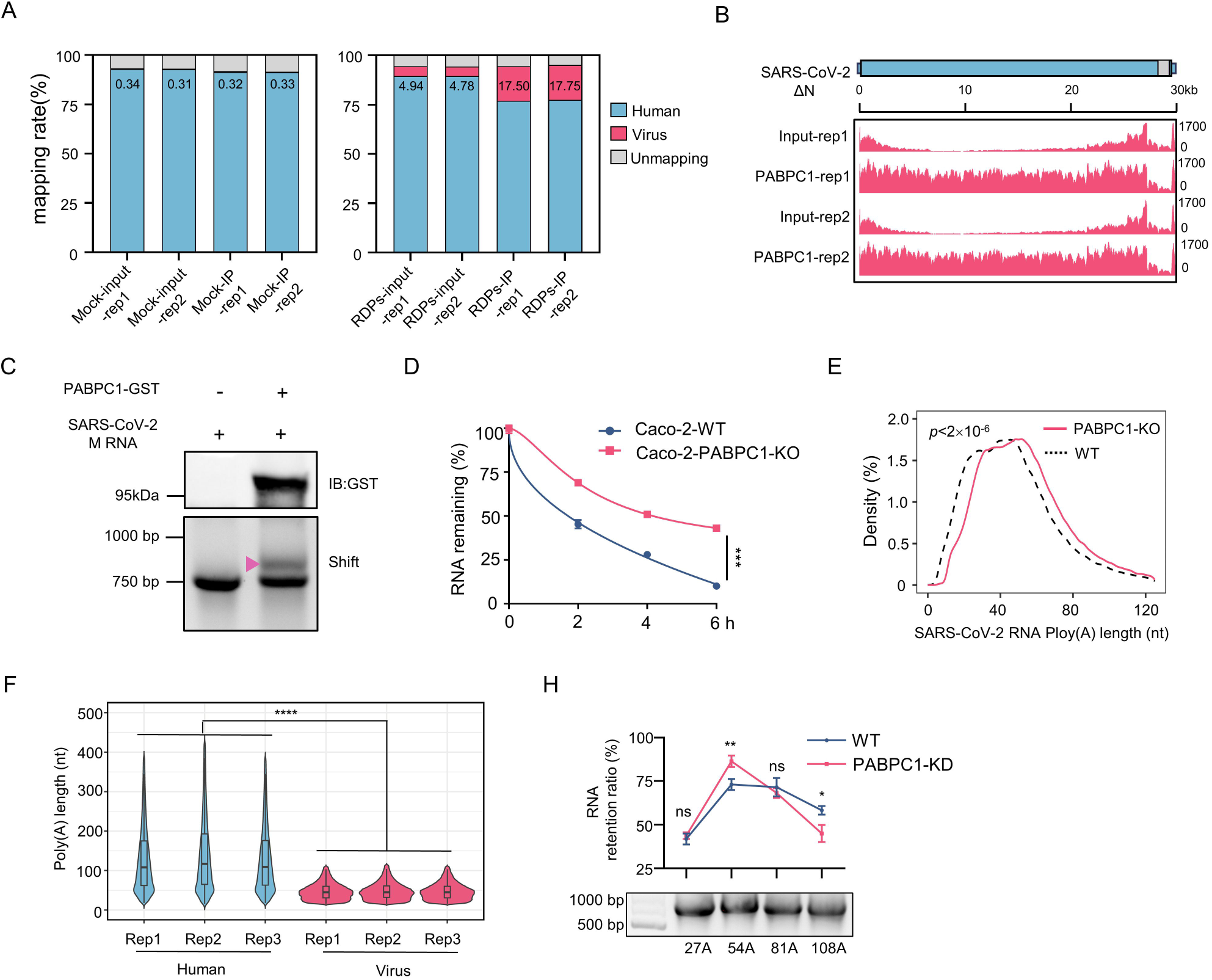
PABPC1 directly binds coronavirus RNA and selectively accelerates the decay of short poly(A)-tailed viral transcripts. (A) RIP-seq analysis of SARS-CoV-2 and human genome mapping rates in input and PABPC1 immunoprecipitated (IP) samples from Caco-2-N cells, with or without SARS-CoV-2 RDPs infection. (B) Integrative Genomics Viewer (IGV) tracks showing read distributions from RIP-seq and input samples along the SARS-CoV-2 positive-sense RNA genome. (C) EMSA analysis of SARS-CoV-2 M RNA binding to purified GST–PABPC1 protein. (D) Stability analysis of RNA of SARS-CoV-2 M gene in Caco-2 WT or PABPC1^-/-^cells with treatment of actinomycin D (ActD) for another 0, 2, 4, and 6 h. (E) Tail-length distribution of viral mRNAs in Caco-2 WT or PABPC1^-/-^cells with infection of SARS-CoV-2. (F) Violin plots showing the poly(A) tail length distributions of human and SARS-CoV-2 mRNAs. (G) Line graph showing RNA retention measured by qPCR in cells with or without PABPC1 knockdown after transfection of luciferase gene containing different poly(A) tail lengths. Data are representative of three independent experiments and were analyzed by two-tailed unpaired t test. Graphs show the mean ± SD (n = 3) derived from three independent experiments. NS, not significant for *P* > 0.05, *\*P* < 0.05, *\*\*P* < 0.01, *\*\*\*P* < 0.001.

### PABPC1-EXD2 interaction drives the degradation of viral RNA

To investigate the mechanism underlying PABPC1’s antiviral activity, we performed IP-mass spectrometry in SARS-CoV-2 RDPs-infected Caco-2-N cells to identify PABPC1-associated proteins (**table S2**). GO and KEGG analyses showed that these interactors were broadly enriched in RNA regulatory pathways, including mRNA surveillance, nucleocytoplasmic transport, and RNA polymerase II–related processes, as well as spliceosomal and preribosomal complexes (**fig. S7A**). Applying stringent filters (Fold change > 2; *P*< 0.05), we identified six high-confidence RNA metabolism factors—YTHDC2, ADAR, ZFC3H1, EXD2, CPSF6, and EDC3—implicated in m A-dependent regulation, A-to-I editing, nuclear RNA surveillance, exonucleolytic decay, 3’-end processing, and decapping-mediated degradation (**Fig. 6A**). To determine which PABPC1-interacting proteins functionally influence SARS-CoV-2 replication, we synthesized siRNAs targeting the six RNA metabolism factors identified by IP-MS. All siRNAs achieved efficient knockdown in Caco-2-N cells, validating their suitability for downstream assays (**fig. S7B**). We next infected Caco-2-N cells with SARS-CoV-2 RDPs to evaluate the functional relevance of each factor. Among the candidates, EXD2, a 3’-5’ exonuclease, emerged as a prominent interactor in regulating of viral replication, suggesting that PABPC1 may recruit EXD2 to promote viral RNA degradation (**Fig. 6B**). We next focused on EXD2 as a key PABPC1-interacting factor. Co-immunoprecipitation assays confirmed the interaction between PABPC1 and EXD2 (**Fig. 6C**), and in vitro GST pulldown experiments demonstrated that this interaction is direct (**fig. S7C, Fig. 6D**). Confocal microscopy further revealed clear cytoplasmic colocalization of the two proteins (**Fig. 6E**). Building on previous studies, we explored the structural basis for PABPC1-EXD2 interaction. PABPC1 is a 636-residue basic protein comprising four N-terminal RNA-recognition motifs (RRM1-4), a proline-rich intrinsically disordered linker of ∼170 residues, and a C-terminal PABC domain^8,39,40^. EXD2 is a 621-residue mitochondria-associated DEDDh-family nuclease with an N-terminal transmembrane helix that anchors it to the outer mitochondrial membrane, a central metal-dependent 3′-5′ exonuclease domain containing canonical catalytic motifs, and a C-terminal HNH-like module implicated in substrate recognition^21^. AlphaFold3 structural predictions suggested that PABPC1 engages EXD2 through contacts between its RRM1-2 domains and the region spanning EXD2’s 3’-5’ exonuclease and HNH domains (**fig. S7D**). To experimentally validate this interface, we generated a series of PABPC1 and EXD2 truncation mutants (**Fig. 6F-G**). Domain-mapping Co-IP assays showed that the N-terminal 1-89 aa of PABPC1 interacts specifically with the 296-408 aa region of EXD2, confirming the predicted binding interface (**Fig. 6H-I**).

**Fig. 6.**
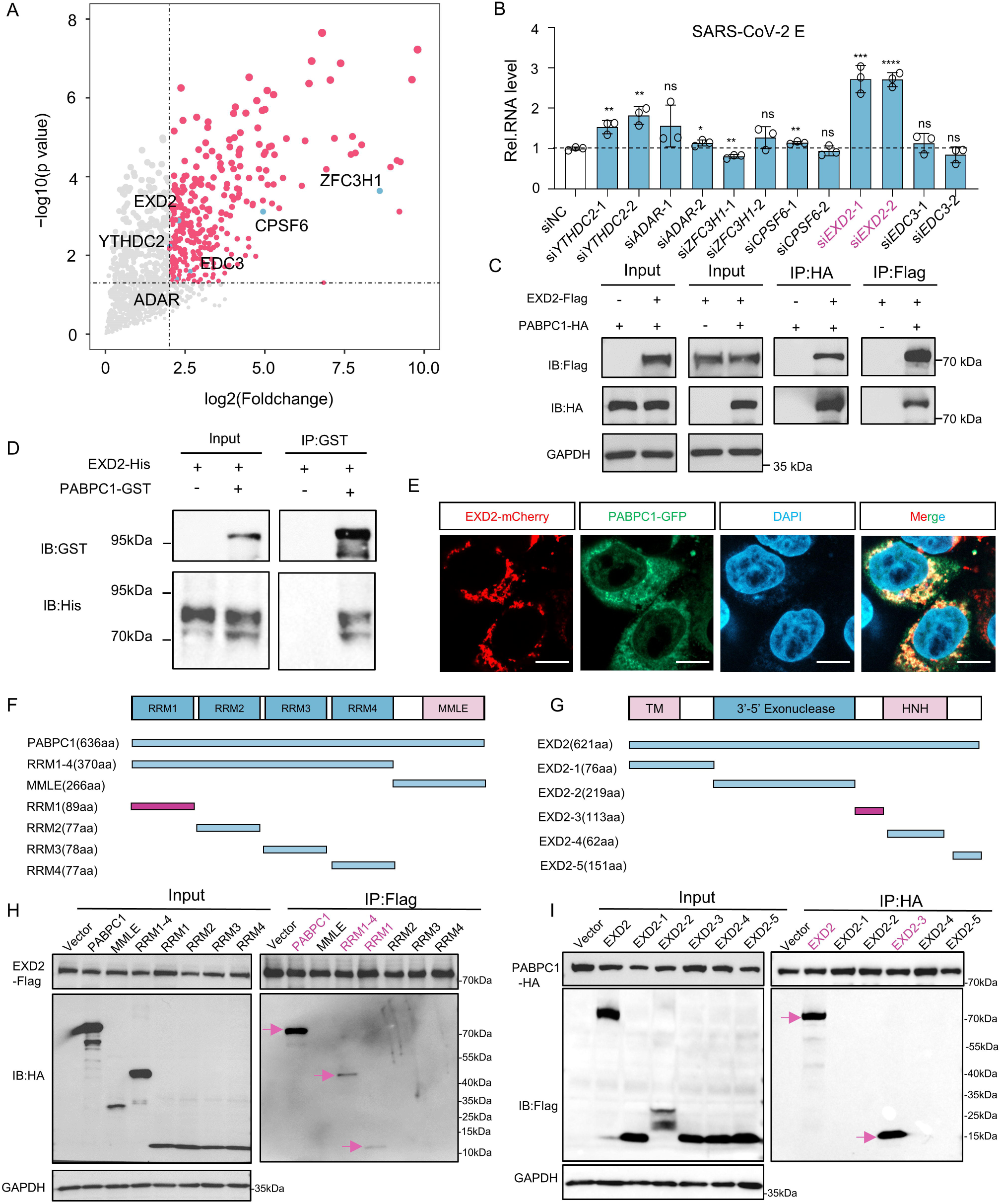
PABPC1-EXD2 interaction drives the degradation of viral RNA. (A) Scatter plot showing identification of PABPC1-interacting proteins by IP-MS, with six RNA metabolism factors highlighted. (B) qPCR analysis of SARS-CoV-2 E gene RNA levels following knockdown of PABPC1-interacting proteins and infection with SARS-CoV-2 RDPs. (C) Co-immunoprecipitation analysis of PABPC1-HA and EXD2-Flag co-expressed in HEK293T cells. (D) GST pulldown analysis of bacterially expressed and purified PABPC1-GST and EXD2-His proteins. (E) Fluorescence images showing co-localization of PABPC1-GFP and EXD2-mCherry. Scale bar, 100 µm. Schematic representation of PABPC1 (F) and EXD2 (G) truncation constructs. Co-immunoprecipitation analysis of PABPC1 truncation fragments co-expressed with full-length EXD2 in HEK293T cells (H), and of EXD2 truncation fragments co-expressed with full-length PABPC1 (I). Data are representative of three independent experiments and were analyzed by two-tailed unpaired t test. Graphs show the mean ± SD (n = 3) derived from three independent experiments. NS, not significant for *P* > 0.05, *\*P* < 0.05, *\*\*P* < 0.01, *\*\*\*P* < 0.001.

Functional characterization of EXD2 revealed that its loss markedly promoted SARS-CoV-2 replication. EXD2 knockdown significantly enhanced both SARS-CoV-2 RDPs and RDPs propagation, as shown by elevated luminescence, increased GFP fluorescence, and time-dependent upregulation of viral E and ORF1ab transcripts (**Fig. 7A-C**). Conversely, SARS-CoV-2-Luc reporter assays demonstrated that EXD2 overexpression potently suppressed reporter activity in a dose-dependent manner (**Fig. 7D**). Truncation analysis further identified the EXD2-2 fragment, corresponding to its 3’-5’ exonuclease domain, as sufficient to mediate this antiviral effect (**Fig. 7E**). To directly assess its biochemical activity, full-length EXD2 and an exonuclease-deficient ΔEXD2 variant were bacterially expressed and purified from *E. coli* BL21 (**Fig. 7F**). *In vitro* RNA degradation assays demonstrated that only full-length EXD2, but not ΔEXD2, efficiently degraded viral RNA (**Fig. 7G**), establishing the 3’-5’ exonuclease domain as the critical module required for EXD2-mediated restriction of SARS-CoV-2. These results indicate that the interaction between PABPC1 and EXD2 is critical for the degradation of viral RNA.

**Fig. 7.**
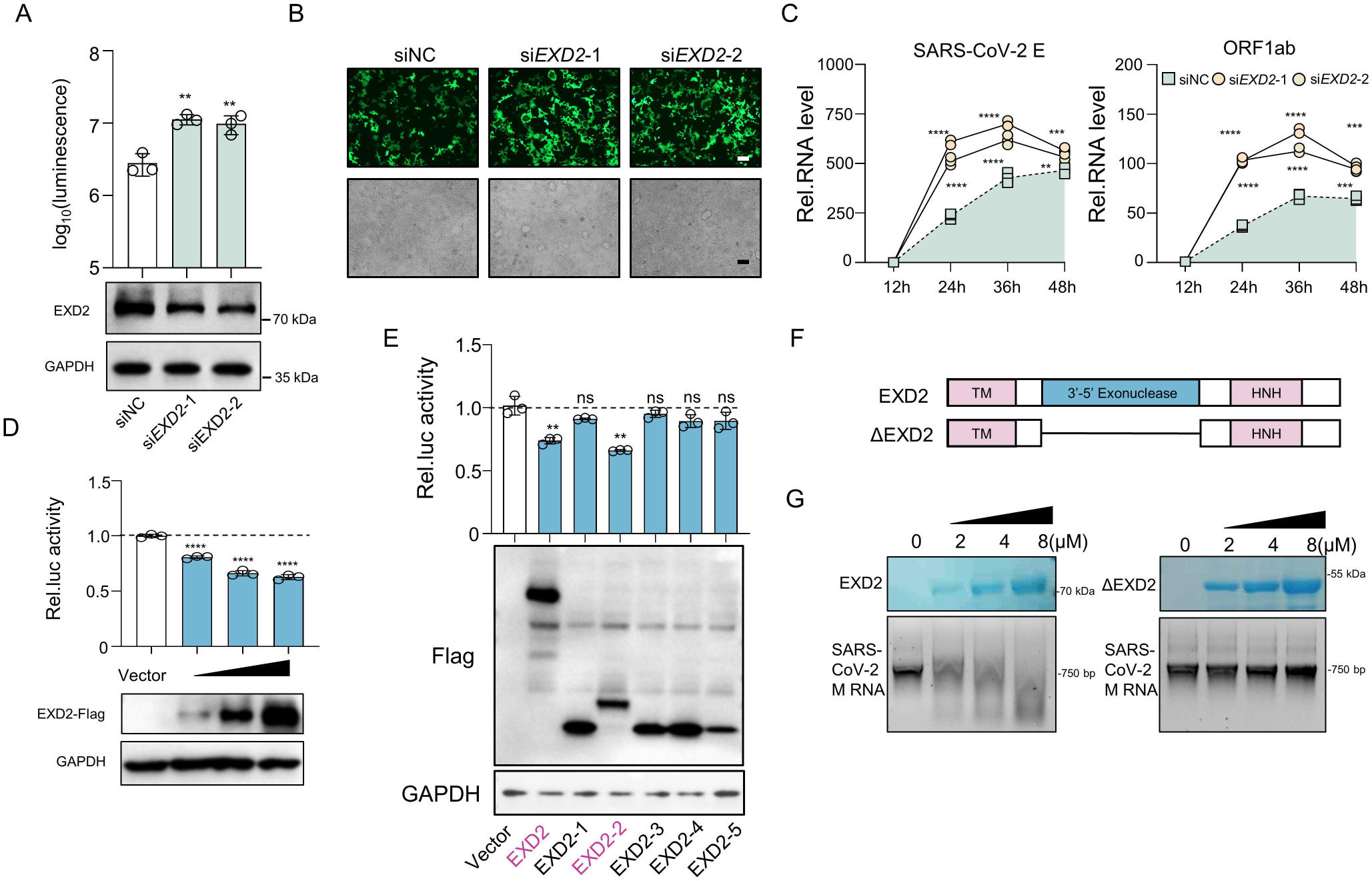
EXD2 mediates antiviral activity through 3’-5’ exonuclease activity. (A-C) Kinetics of HiBiT luminescence, GFP fluorescence, and qPCR analysis of SARS-CoV-2 E and ORF1ab RNA levels in Caco-2-N cells following EXD2 knockdown and infection with SARS-CoV-2 RDPs. (D) Relative luciferase activity of the SARS-CoV-2-Luc reporter in cells transfected with 100 ng, 200 ng, or 500 ng of EXD2-Flag. (E) Relative luciferase activity of the SARS-CoV-2-Luc reporter in cells transfected with 500 ng of EXD2 truncation fragments. (F) Schematic representation of EXD2 and ΔEXD2. (G) Agarose gel electrophoresis analysis of SARS-CoV-2 M RNA following incubation with full-length EXD2 or the exonuclease-deficient ΔEXD2 protein. Data are representative of three independent experiments and were analyzed by two-tailed unpaired t test. Graphs show the mean ± SD (n = 3) derived from three independent experiments. NS, not significant for *P* > 0.05, *\*P* < 0.05, *\*\*P* < 0.01, *\*\*\*P* < 0.001.

### The fusion protein of the key functional domains of PABPC1 and EXD2 inhibits coronavirus replication as a novel nucleic acid therapeutic

Based on our previous findings, Caco-2-N cells were transfected with plasmids expressing PABPC1 and EXD2. Individually, neither PABPC1 nor EXD2 significantly inhibited SARS-CoV-2 RDPs replication (**fig. S8A-F**). Intriguingly, co-expression of PABPC1 and EXD2 markedly suppressed viral replication in a dose-dependent manner, indicating a synergistic effect (**fig. S8G-I, Fig. 8A**). Leveraging the functional domains responsible for RNA binding (PABPC1 RRM1-4) and exonuclease activity (EXD2 3’-5’ exonuclease), we engineered a fusion protein connected by a flexible linker, with an estimated molecular weight of ∼55 kDa (**Fig. 8B**). Alphafold3 structural predictions confirmed proper domain organization (**Fig. 8C**). Functional evaluation using the SARS-CoV-2-Luc reporter system demonstrated enhanced, dose-dependent viral inhibition (**Fig. 8D**). Similarly, transfection of Caco-2-N cells with the fusion construct significantly reduced replication of SARS-CoV-2 RDPs (**Fig. 8E-G**). To explore the therapeutic potential of the fusion protein, we i*n vitro*-transcribed its RNA and encapsulated it using our lipid nanoparticle (LNP) system^41–43^, generating a novel nucleic acid-based antiviral agent.

**Fig. 8.**
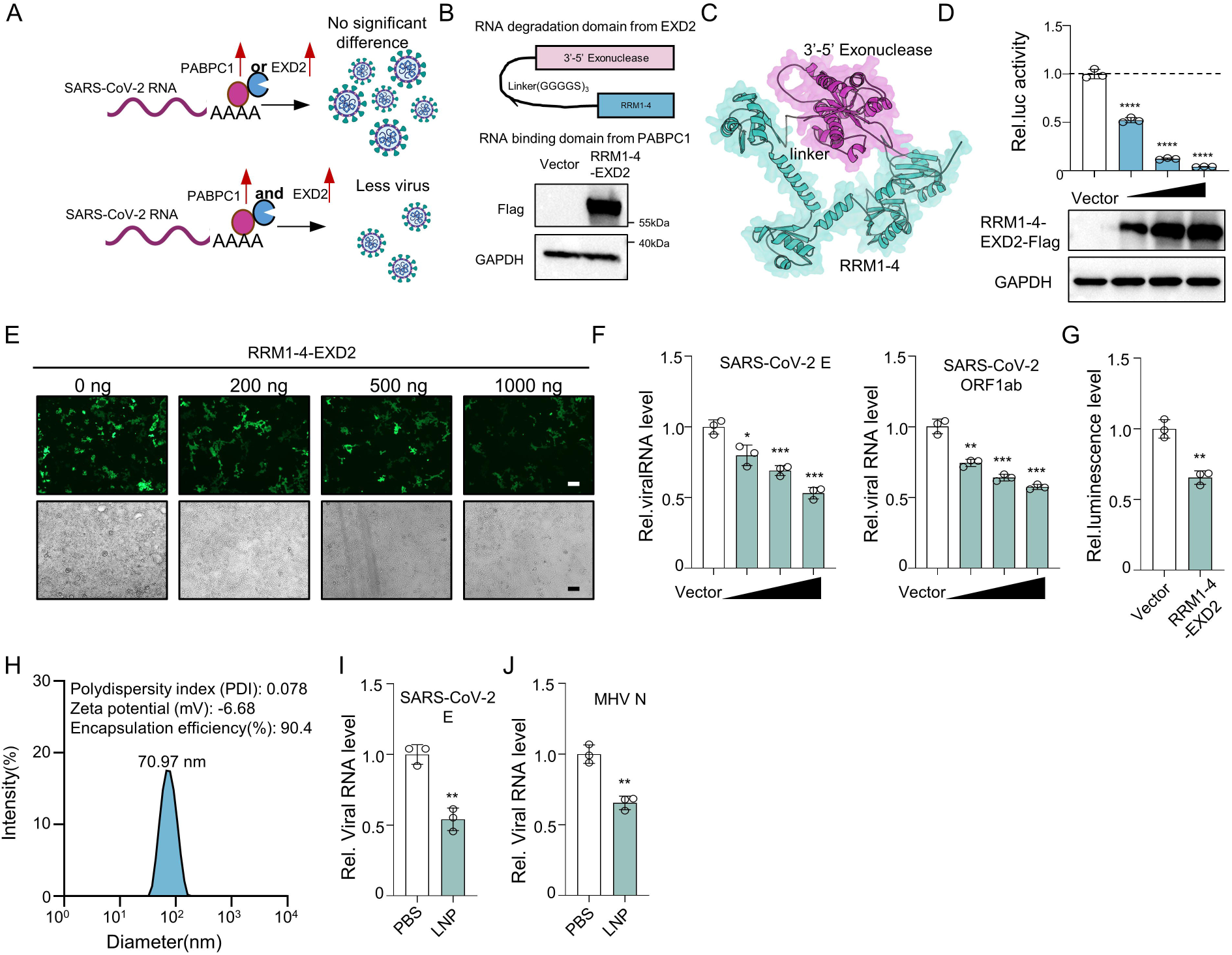
The fusion protein of the key functional domains of PABPC1 and EXD2 inhibits SARS-CoV-2 and MHV replication as a novel nucleic acid therapeutic. (A) Schematic representation illustrating that co-expression of PABPC1 and EXD2 suppresses SARS-CoV-2 RDPs replication. (B) Schematic representation and Western blot analysis of the PABPC1–EXD2 fusion protein. (C) AlphaFold3-predicted structure of the PABPC1–EXD2 fusion protein. (D) Relative luciferase activity of the SARS-CoV-2-Luc reporter in cells transfected with 100 ng, 200 ng, or 500 ng of RRM1-4-EXD2-Flag. (E-G) GFP fluorescence, qPCR analysis of SARS-CoV-2 E and ORF1ab RNA levels, and kinetics of HiBiT luminescence in Caco-2-N cells transfected with RRM1-4–EXD2-Flag and infected with SARS-CoV-2 RDPs. (H) Characterization of lipid nanoparticle (LNP)-encapsulated fusion mRNA. (I) qPCR analysis of SARS-CoV-2 E RNA levels in Caco-2-N cells following LNP treatment and infection with SARS-CoV-2 RDPs. (J) qPCR analysis of MHV N RNA levels in Neruo2a cells following LNP treatment and infection with MHV. Data are representative of three independent experiments and were analyzed by two-tailed unpaired t test. Graphs show the mean ± SD (n = 3 in D, F-G, I-J) derived from three independent experiments. NS, not significant for *P* > 0.05, *\*P* < 0.05, *\*\*P* < 0.01, *\*\*\*P* < 0.001.

Characterization of the lipid nanoparticles revealed a hydrodynamic diameter of 70.97 nm and a polydispersity index (PDI) of 0.078, indicating a uniform particle population. The surface charge was nearly neutral. Furthermore, the encapsulation efficiency of mRNA was determined to be >90%. Collectively, these data indicate that the LNP possessed good physical properties (**Fig. 8H**). In cell culture, this LNP markedly inhibited replication of both SARS-CoV-2 RDPs and MHV (**Fig. 8I-J**).

### Inflammatory stimuli upregulate PABPC1 and EXD2 Expression

Viral infection and host defense are in a constant arms race. We further collected RNA from uninfected and SARS-CoV-2-infected cells and found the mRNA levels PABPC1 and EXD2 were significantly upregulated (**Fig. 9A**), consistent with our previous RNA-seq data (HRA006231)^31^ (**Fig. 9B**). Analysis of the GSE217504 dataset also revealed dynamic upregulation of PABPC1 and EXD2 upon SARS-CoV-2 infection^44^ (**Fig. 9C**). Moreover, human infection datasets, including CRA002390, further confirmed the elevated expression of both factors in patients (**Fig. 9D**), indicating that PABPC1 and EXD2 are induced as part of the host response to viral infection. To investigate the *in vivo* dynamics of PABPC1 and EXD2 during SARS-CoV-2 infection, we utilized K18-hACE2 C57/BL6J mice^45^. Mice were intranasally infected with 2,500 PFU of SARS-CoV-2 BA.2 and sacrificed on days 3, 5, and 7 post-infection for lung collection (**Fig. 9E**). Mild body weight loss was observed over time (**Fig. 9F**). qPCR analysis revealed consistently high viral loads in lung tissues across all three time points (**Fig. 9G**). RNA-seq analysis showed high reproducibility among replicates, as confirmed by Pearson correlation and PCA analyses (**fig. S9A-B)**. GSEA revealed that inflammatory pathways, including Chemokine signaling, Toll-like receptor signaling, and NOD-like receptor signaling, peaked on day 5 and declined by day 7, indicating a transient inflammatory surge (**Fig. 9H**). Heatmap visualization of inflammatory cytokines supported this temporal pattern (**fig. S9C)**. Histopathological analysis showed more severe lung injury on days 5 and 7 compared to day 3 (**Fig. 9I**). qPCR confirmed elevated Il-1β, Il-6, and Il-8 expression following the same trend (**Fig. 9J**). Notably, the mRNA levels of *Pabpc1* and *Exd2* were also increased on days 5 and 7, suggesting potential regulation by inflammation (**Fig. 9K**). To directly test this hypothesis, we stimulated mice with LPS for 24 h and analyzed lung RNA by RNA-seq (data from ref^33^) (**Fig. 9L**). GSEA revealed significant activation of multiple inflammatory pathways, including Cytokine-cytokine receptor interaction, NF-κB signaling, NOD-like receptor signaling, TNF signaling, and Toll-like receptor signaling (**Fig. 9M**). Volcano plot analysis indicated upregulation of both *Pabpc1* and *Exd2* (**Fig. 9N**). In A549 cells, treatment with IL-1β or TNFα similarly induced robust upregulation of *PABPC1* and *EXD2*, alongside canonical inflammatory markers *IL6*, *IL8*, and *TNF*α, as confirmed by qPCR (**fig. S9D-G**). Western blotting and densitometric quantification further demonstrated increased protein levels of PABPC1 and EXD2 upon inflammatory cytokine stimulation (**fig. S9H**), indicating that their expression is responsive to host inflammatory signals.

**Fig. 9.**
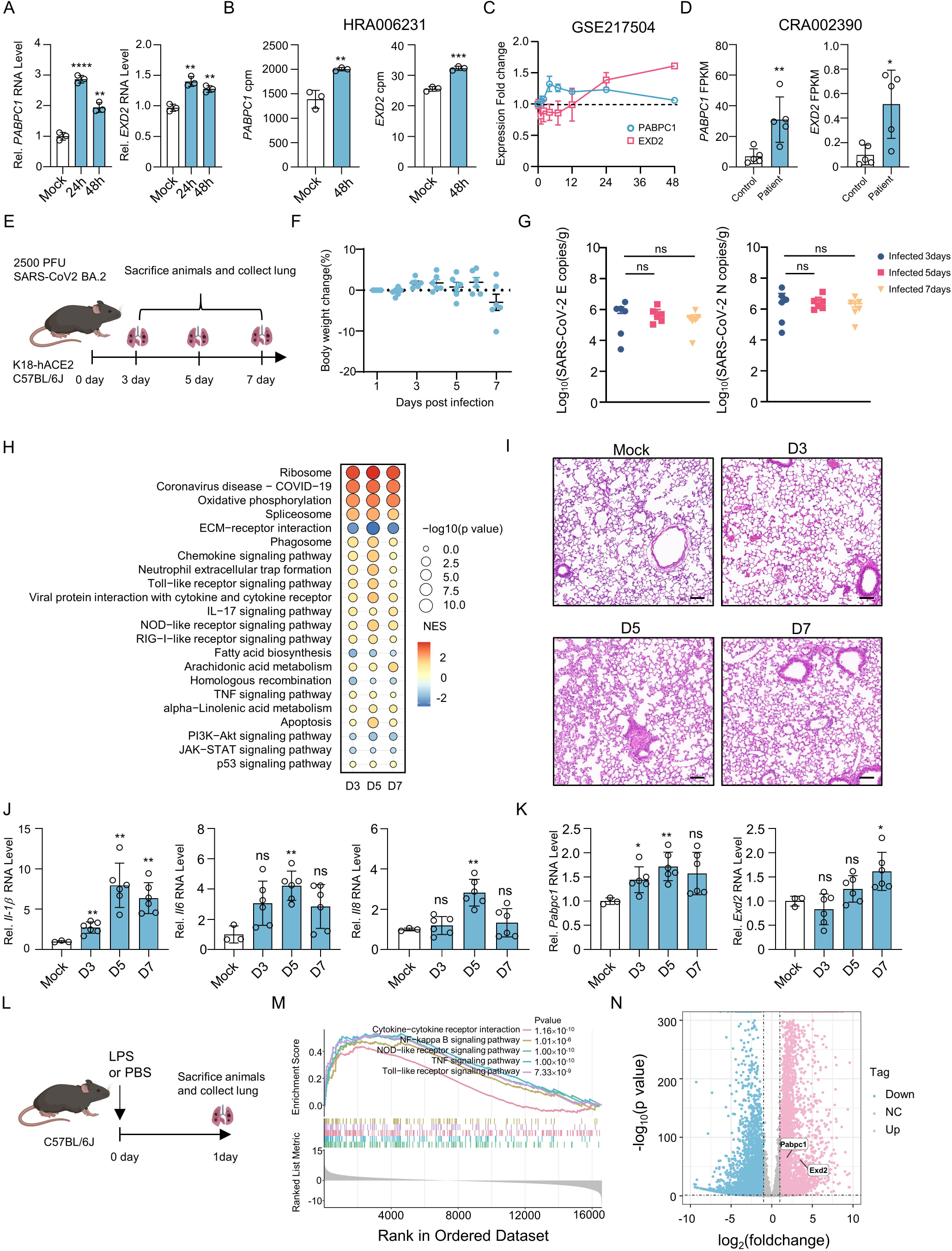
Inflammatory stimuli upregulate PABPC1 and EXD2 Expression (A) qPCR analysis of PABPC1 and EXD2 mRNA levels in Caco-2 cells following infection with SARS-CoV-2. (B) RNA-seq analysis of PABPC1 and EXD2 counts per million (CPM) in Caco-2 cells following SARS-CoV-2 infection. (C) RNA-seq analysis of fold changes in PABPC1 and EXD2 expression in Caco-2 cells at 0, 1, 4, 7, 12, 24, and 48 h post SARS-CoV-2 infection. (D) RNA-seq analysis of FPKM values for PABPC1 and EXD2 expression in SARS-CoV-2-infected patients and healthy controls. (E) Schematic representation of K18-hACE2 mice intranasally infected with SARS-CoV-2 BA.2 and lung collection at 3, 5, and 7 days post-infection. (F) Line graph showing body weight changes in K18-hACE2 mice, n=6. (G) qPCR analysis of SARS-CoV-2 E and N RNA levels in the lung of K18-hACE2 mice, n=6. (H) GSEA enrichment analysis of pathways in lungs of K18-hACE2 mice at 3, 5, and 7 days post SARS-CoV-2 BA.2 infection. (I) Representative H&E-stained lung sections from uninfected mice and mice at 3, 5, and 7 days post-infection, n=6. Scale bar, 200 µm. (J) qPCR analysis of *IL-1*β, *IL-6* and *IL-8* mRNA levels in lungs of K18-hACE2 mice at 3, 5, and 7 days post SARS-CoV-2 BA.2 infection. (K) qPCR analysis of PABPC1 and EXD2 mRNA levels in lungs of K18-hACE2 mice at 3, 5, and 7 days post SARS-CoV-2 BA.2 infection. (L) Schematic representation of C57BL/6J mice sacrificed 1 day after intraperitoneal injection of LPS for lung collection. (M) GSEA enrichment analysis of signaling pathways in the lungs of C57BL/6J mice following intraperitoneal injection of LPS. (N) Volcano plot analysis of differential gene expression in the lungs of C57BL/6J mice before and after LPS treatment (fold change > 2, *P* < 0.05). Data are representative of three independent experiments and were analyzed by two-tailed unpaired t test. Graphs show the mean ± SD (n = 3 in A-C; n = 5 in F-G, J-K) derived from three independent experiments. NS, not significant for *P* > 0.05, *\*P* < 0.05, *\*\*P* < 0.01, *\*\*\*P* < 0.001.

## Discussion

Viral RNA hijacks host RBPs that govern various stages of mRNA metabolism to orchestrate translation and replication under the hostile intracellular environment. Although numerous studies have characterized coronavirus RBPs, discrepancies often arise due to differences in infection methods and viral titers^1,2,22–26^. In this study, we defined 94 high-confidence RBPs as those consistently identified in more than four out of nine independent datasets (**Fig. 1A-D**). To further refine the core set of RBPs, we intersected these with RBPs reported for other RNA viruses. Consistent with the observations of Flynn et al.^26^, whose viral RNA interactome exhibited a pronounced enrichment for antiviral factors, our analysis also revealed a similar bias towards host antiviral RBPs (**Fig. 1E-H**).

Our findings establish PABPC1 as a key host factor with broad-spectrum antiviral activity against coronaviruses. While PABPC1 is a well-characterized poly(A)-binding protein, its functional impact varies across viral infections^12,14–16,46–50^. Our results extend these findings, demonstrating that PABPC1 exerts potent antiviral activity across multiple CoVs, including SARS-CoV-2, HCoV-OC43, and MHV (**Fig. 5**). This suggests that PABPC1-mediated restriction may represent a conserved host defense mechanism targeting coronavirus RNA.

While characterizing PABPC1 activity, we identified two AML-targeting small-molecule inhibitors that competitively disrupt PABPC1 binding to poly(A) RNA^37^. Functional assays in our coronavirus infection models revealed that both compounds markedly enhanced SARS-CoV-2 and HCoV-OC43 replication, consistent with their interference with PABPC1-mediated antiviral defense (**Fig. 3, fig. S5**). These findings extend our understanding of PABPC1 as a critical restriction factor and highlight that therapeutic agents designed to target PABPC1-dependent processes in cancer may inadvertently potentiate viral infection (**Fig. 10A**).

**Fig. 10.**
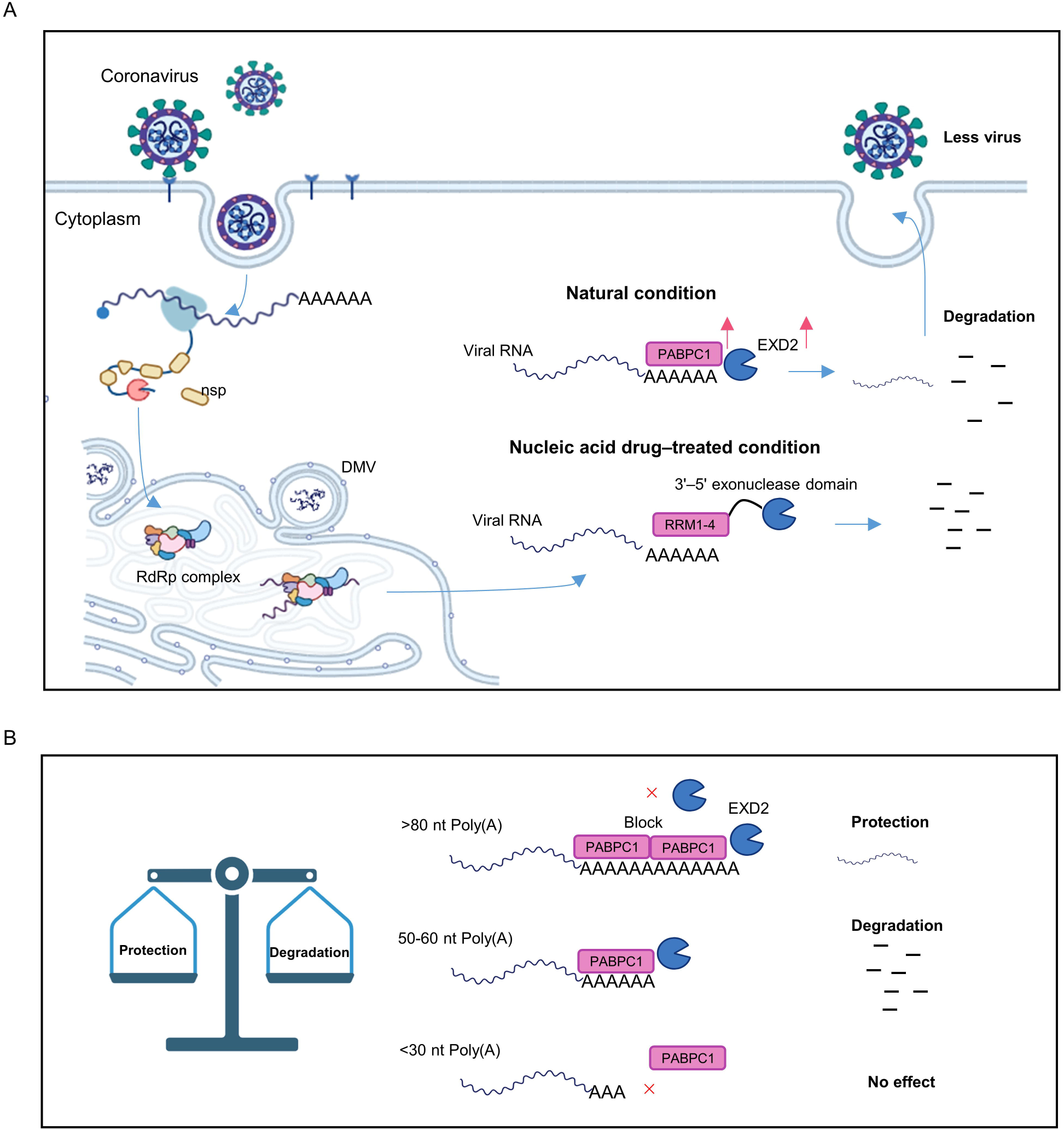
Poly(A)-tail-length–dependent recognition of viral RNA by PABPC1 enables selective recruitment of EXD2 and targeted degradation of short-tailed coronavirus transcripts (A) Under physiological conditions, PABPC1 and EXD2 are upregulated upon viral infection through inflammatory signaling and cooperate to degrade viral RNA. Our engineered fusion-protein nucleic acid therapeutic harnesses this endogenous antiviral mechanism, thereby achieving superior antiviral efficacy. (B) Our data suggest that tail length critically biases this regulatory outcome: short poly(A) tails are insufficient for effective PABPC1 binding, whereas intermediate-length tails (50–60 nt) are preferentially targeted by PABPC1 for degradation. As the poly(A) tails lengthen, this degradative effect is attenuated due to PABPC1-mediated interactions, and the protective and stabilizing functions of PABPC1 become more prominent.

In SARS-CoV-2–infected cells, we observed that PABPC1 selectively associates with viral RNAs (**Fig. 4A**), consistent with prior findings in IAV^13^. This preferential binding likely reflects the distinct poly(A)-tail length distributions between viral and host transcripts, as SARS-CoV-2 genomic and subgenomic RNAs harbor relatively short and narrowly distributed poly(A) tails of ∼50–60 nt (**Fig. 4G**). Functionally, PABPC1 depletion markedly increased the half-life of viral RNAs (**Fig. 4E**), with the strongest effect observed for RNAs bearing 50–60 nt poly(A) tails (**Fig. 4H**). PABPC1 has been shown to recruit diverse effector proteins and, through binding to poly(A) tails, modulate mRNA stability by either promoting degradation or enhancing transcript protection^10,51–57^. Our data suggest that tail length critically biases this regulatory outcome: short poly(A) tails are insufficient for effective PABPC1 binding, whereas intermediate-length tails (50–60 nt) are preferentially targeted by PABPC1 for degradation. As the poly(A) tails lengthen, this degradative effect is attenuated due to PABPC1-mediated interactions^8^, and the protective and stabilizing functions of PABPC1 become more prominent (**Fig. 10B**).

We subsequently proposed a novel PABPC1-mediated RNA degradation mechanism that relies on EXD2. EXD2 is a mitochondria-associated DEDDh-family nuclease^21^, and here we report for the first time a direct interaction between the RRM1 domain of PABPC1 and the 296–408 region of EXD2 (**Fig. 6F-I**). Our data reveal a mechanistic basis for PABPC1’s antiviral activity, showing that PABPC1 preferentially binds viral poly(A) tails and subsequently recruits EXD2 to mediate targeted degradation of viral RNA. Building on these insights, we engineered a fusion protein combining the functional domains of PABPC1 (RRM1-4) and EXD2 (3’-5’ exonuclease), which demonstrated potent antiviral effects *in vitro* (**Fig. 8A-G**). To further extend the translational potential of this approach, we employed a lipid nanoparticle (LNP) delivery system^43^ for the fusion protein-encoding RNA, achieving robust antiviral efficacy in both cultured cells and mouse models (**Fig. 8H-K**). These findings provide a proof-of-concept for leveraging host RNA-binding and nuclease activities as a modular antiviral strategy (**Fig. 10A**).

Viruses and their hosts are constantly engaged in a dynamic antagonistic interplay^7,58^. Our results indicate that following viral infection, both PABPC1 and EXD2 are upregulated in response to inflammatory signals (**Fig. 9**), suggesting a broad host-mediated antiviral defense mechanism. This observation provides new insights into the role of host immunity in controlling viral replication.

Collectively, we reveal a PABPC1–EXD2–dependent antiviral mechanism whereby PABPC1 preferentially binds viral poly(A) tails and recruits EXD2 to promote RNA degradation. Based on this we developed an effective LNP-delivered therapeutic strategy. Targeting host factors to counteract viral replication represents a promising avenue for antiviral intervention and opens up a new frontier for mechanistic and drug discovery studies from a host-centered perspective.

## Materials and Methods

### Plasmid, viruses, cells, and reagents

The full-length PABPC1 gene was cloned into the pCAGGS-HA vector, and EXD2 was cloned into pCMV-Flag. Both vectors were maintained in our laboratory. For prokaryotic expression, the corresponding genes were subcloned into pET28a and pGEX-6P-1 vectors, which are also preserved in our laboratory. The SARS-CoV-2 luciferase reporter system was constructed by replacing the original pGL3-promoter UTR regions with the 5’ and 3’ UTRs of the SARS-CoV-2 wild-type strain (IVCAS 6.7512). The lentiCRISPR-v2 plasmid (Addgene) was used for knockout experiments. Lentiviral packaging plasmids, including the lentiviral backbone (1 µg), psPAX2 (0.5 µg), and pMD2.G, were all previously maintained our laboratory.

The SARS-CoV-2 wild-type strain (IVCAS 6.7512), Beta variant (B.1.351; NPRC2.062100001), and Omicron variant (BA.2) were obtained from the National Virus Resource, Wuhan Institute of Virology, Chinese Academy of Sciences.

Caco-2, A549, HEK293T, and Huh7 cells were maintained in Dulbecco’s modified Eagle’s medium (DMEM) supplemented with 10% fetal bovine serum, 100 U/mL penicillin, and 100 µg/mL streptomycin at 37 °C in a 5% CO incubator.

Plasmids were transfected using Lipofectamine 3000 (Invitrogen) or Neofect DNA transfection reagent (Neofect, TF201201) according to the manufacturers’ protocols. siRNAs (RiboBio, Guangzhou, China) were delivered using RNAiMAX (Invitrogen). Ruxolitinib and actinomycin D were purchased from MedChemExpress (MCE). For stimulation experiments, recombinant human IL-1β (Peprotech, 200-01B) and TNF-α (Peprotech, 300-01A) were applied at 10 ng/mL, and lipopolysaccharide (LPS; Sigma-Aldrich, L2630) was administered at 25 mg/kg via intraperitoneal injection in mice.

### Mice and ethics statement

All animal experiments and methods were performed following the relevant approved guidelines and regulations, as well as under the approval of the Ethics Committee of college of life sciences at Wuhan University. *Pabpc1*^-/-^ C57BL/6J mice were obtained from Gempharmatech Co., Ltd. (Nanjing, China) and maintained under specific pathogen-free (SPF) conditions. Genotyping of Pabpc1 mutant mice was performed using the following primers: F1, TTTCGAGTACCCTCCTGGAG; R1, CGTGTCTTTAATCCCAACACTCG (WT: 3830 bp; Targeted: 334 bp); F2, TGAAGTTCCTGGGATTGGGTG; R2, GGCCTGGCTGTTCTAACCTATGTAG (WT: 430 bp; Targeted: 0 bp). B6/JGpt-H11^em1Cin(K^^18-ACE2)^/Gpt mice (K18-hACE2 KI mice, heterozygote, #T037657) were purchased from GemPharmatech Co. Ltd^45,59^. The Sftpc-N-hACE2 mice were previously generated in our laboratory, with stable insertion of the Sftpc lung-specific promoter driving expression of SARS-CoV-2 N and human ACE2 genes^38^. The Animal Care and Use Committee (IACUC) of the Animal Experimentation Center, Wuhan University, reviewed and approved the submitted Animal Use Protocols (AUPs) in accordance with relevant animal welfare and ethical guidelines, with approval numbers WP20220044 and WP20240486.

### SARS-CoV-2 live virus infection assay

All SARS-CoV-2 live virus-related experiments were approved by the Level 3 Biosafety Committee (ABSL-3) of Wuhan University. All experiments involving SARS-CoV-2 were performed in the BSL-3 and ABSL-3 facility of Wuhan University.

### SARS-CoV-2 **Δ**N-GFP-HiBiT replicon delivery particles

SARS-CoV-2 ΔN-GFP-HiBiT replication-defective particles (RDPs) were constructed as previously described^30^. To determine viral infectivity and titer, the 50% tissue culture infective dose (TCID) endpoint assay was performed. Caco-2-N cells were seeded into 96-well plates and cultured to full confluence. Viral stocks were serially diluted 10-fold in serum-free medium (10 ¹ to 10) with eight replicates per dilution. After removing the culture medium, diluted viruses were added to the cells. Following 48 h of incubation, fluorescent cells in each well were counted under a fluorescence microscope. TCID values were calculated using the Reed-Muench method.

### Construction of knockout cell line by CRISPR/Cas9

The PABPC1-targeting gRNA was designed as follows: gRNA-F: CACCGAAAGTAATGACTGATGAAAG, gRNA-R: AAACCTTTCATCAGTCATTACTTTC, and cloned into the lentiCRISPR-v2 plasmid (Addgene). Lentiviral particles were generated by co-transfecting the lentiviral backbone (1 µg), psPAX2 (0.5 µg), and pMD2.G (0.5 µg) into HEK293T cells seeded in 6-well plates using Neofect. After 48 h, viral supernatants were collected, filtered through 0.45 µm membranes, and used to infect target cells in the presence of polybrene (8 µg/mL). Cells were infected twice to enhance transduction efficiency. Puromycin selection was applied to enrich for positive cells, which were then seeded at one cell per well in 96-well plates. After two weeks of growth under selection, single-cell clones were expanded, and knockout efficiency was confirmed by immunoblotting, genomic PCR, and sequencing.

### Dual-luciferase assay

HEK293T cells were seeded in 24-well plates and transfected with plasmid combinations, including 100 ng of a SARS-CoV-2–luciferase reporter plasmid and 20 ng of pRL-TK Renilla luciferase plasmid as an internal control. After transfection, cells were lysed with passive lysis buffer, and luciferase activities were measured using the Dual-Luciferase Reporter Assay System (E1910, Promega). Firefly luciferase activity (value A) was first measured by adding 20 µL of substrate I to 20 µL of cell lysate per well, followed by measurement of Renilla luciferase activity (value B) with 20 µL of substrate II. Relative reporter expression was calculated as the A/B ratio. All assays were performed in three independent biological replicates.

### Antibodies and immunoblot analysis

The antibodies used were: rabbit anti-PABPC1 (Sino Biological, 105506-T44), rabbit anti-EXD2 (Proteintech, 20138-1-AP), mouse anti-SARS-CoV-2 Nucleocapsid (Sino Biological, 40143-MM05), mouse anti-Flag (Sigma, F3165), mouse anti-GAPDH (Proteintech, 60004-1-Ig), mouse anti-His (Proteintech, HRP-66005), and mouse anti-GST (Proteintech, 10000-0-AP). Cells were washed once with PBS and lysed in RIPA or NP-40 lysis buffer (Beyotime). Protein samples were mixed with 5× SDS loading buffer, boiled for 10 min, resolved by SDS-PAGE, and transferred to nitrocellulose membranes (GE Healthcare). Membranes were blocked with TBST containing 5% non-fat milk or BSA and probed with primary and secondary antibodies.

For co-immunoprecipitation, HEK293T cells were seeded in 6-cm dishes and transfected with 3 μg of empty or expression plasmids. At 36 h post-transfection, cells were lysed in NP-40 buffer supplemented with protease inhibitors (HY-K0010; MCE) at 4 °C for 30 min. Lysates were cleared by centrifugation and incubated with Anti-Flag or Anti-HA magnetic beads (HY-K0207/HY-K0201; MCE) overnight at 4 °C with rotation. Beads were washed six times with TBST/PBST (CR10301S/CR10201S; Monad) before elution and analysis by SDS-PAGE and immunoblotting.

### Immunofluorescence assay

Caco-2 or Huh7 cells were seeded in 48-well plates and infected with SARS-CoV-2 for 24 h. Cells were fixed and inactivated with 4% paraformaldehyde at room temperature for 30 min, permeabilized with 0.2% Triton X-100 for 10 min, and blocked with 1% BSA at 37 °C for 30 min. Cells were then incubated with mouse anti–SARS-CoV-2 Nucleocapsid antibody (Sino Biological, 40143-MM05) diluted in 1% BSA at 37 °C for 1 h, followed by three washes with PBS. Secondary staining was performed using Alexa Fluor 594-conjugated goat anti-mouse IgG (Thermo Fisher Scientific, A11032) diluted in 1% BSA at room temperature for 30 min, followed by PBS washes. Nuclei were counterstained with Hoechst 33342 for 5 min. Fluorescence images were acquired using an inverted fluorescence microscope.

### GST pull-down assay

The pET28a-EXD2-His plasmid was transformed into *E. coli* BL21 competent cells, and a single colony was inoculated into LB medium. When the culture reached an OD of 0.6–0.8, protein expression was induced with 0.4 mM IPTG at 18 °C overnight (∼18 h). Cells were harvested, lysed, and the lysate incubated with Ni-NTA resin (L00250; Genscript) at 4 °C for 4 h. After washing to remove non-specific proteins, His-tagged EXD2 was eluted and quantified by OD. Similarly, pGEX-6P-1-PABPC1-GST plasmids were transformed into *E. coli* BL21, and expression and purification were performed using the GST Fusion Protein Purification Kit (L00207; Genscript). For in vitro interaction assays, His-tagged EXD2 and GST-tagged PABPC1 were mixed in PBS and incubated for 4 h at 4 °C, followed by addition of GST magnetic beads (MA113-25T; ABMagic). After overnight incubation at 4 °C, beads were washed five times with pre-cooled PBST. Bound proteins were eluted by boiling in 1% SDS for 15 min, and samples were analyzed by SDS-PAGE and immunoblotting.

### Confocal microscopy

HeLa cells were seeded onto coverslips and transfected with GFP-PABPC1 and mCherry-EXD2 plasmids for 24 h. Cells were then fixed with 4% paraformaldehyde for 15 min at room temperature, and nuclei were counterstained with DAPI. Confocal images were acquired using a Zeiss LSM 980 laser scanning microscope.

### RNA isolation and qPCR

Total RNA was extracted using TRIzol reagent (Invitrogen) according to the manufacturer’s protocol. cDNA was synthesized using PrimeScript RT Reagent Kit (Takara, RR037A) or NovoScript Plus All-in-one 1st Strand cDNA Synthesis SuperMix (Novoprotein). Quantitative real-time PCR (qPCR) was performed on an ABI 7500 Real-Time PCR System using SYBR Green Master Mix (YEASEN, 11199ES03) or TaqMan Probe Master Mix (YEASEN, 11205ES08). Gene expression levels were normalized to GAPDH and calculated using the ΔΔCt method.

### In vitro transcription assays

RNAs were synthesized in vitro using UTPs (Syngenebio, Nanjing, China) as substrates with the MEGAscript T7 Transcription Kit (Ambion, USA) following the manufacturer’s protocol. For transfection experiments, the transcribed RNAs were enzymatically capped using Vaccinia Capping Enzyme (NEB, M2080) and mRNA Cap 2’-O-Methyltransferase (NEB, M0366) in the presence of GTP and SAM to generate fully capped and methylated transcripts.

### Electrophoretic Mobility Shift Assay (EMSA)

EMSA reactions were assembled on ice in a total volume of 20 µL, containing 50 mM Tris buffer, purified protein, purified RNA, and RNase inhibitor, with DEPC-treated water added to reach the final volume. Reactions were incubated on ice for 30 min, followed by addition of 2× RNA loading buffer (ThermoFisher, R0641). Samples were resolved on a pre-cast TBE agarose gel at 130 V on ice for 50 min, and RNA–protein complexes were visualized using standard nucleic acid staining.

### Formulation of lipid nanoparticle (LNP) therapeutics

The delivery system was prepared by mixing the four lipids ALC-0315, ALC-0159, DSPC, and cholesterol (Sinopeg, China) at a molar ratio of 46.3:1.6:9.4:42.7. The mRNA was diluted in 20 mM sodium acetate buffer at pH 4.0 (Coolaber, China). Subsequently, for nanoparticle formation, the chip was inserted into the iNano™ L+ (Micro & Nano, China), where the organic phase (lipid mixture) and the aqueous phase (diluted mRNA) were introduced via separate syringes. The particles were formed at a total flow rate of 12 mL/min with a flow rate ratio of 1:3 (organic to aqueous). The sodium acetate buffer and ethanol of the resulting sample was replaced with PBS (pH 7.4) using a 100 kDa centrifuge tube (Millipore, United States), followed by filtration through a 0.22 μm membrane. Finally, the purified nanoparticles were characterized by measuring their hydrodynamic diameter and zeta potential using a nanoparticle size analyzer (OMEC, China). The encapsulation efficiency was determined via Ribogreen assay, where the encapsulated RNA concentration was calculated by subtracting the measured free RNA concentration from the total RNA concentration.

### RNA-seq analysis

Total RNA was extracted from the indicated cells using TRIzol reagent (15596026; Ambion) and treated with DNase I (HY-108882; MCE) to remove genomic DNA. RNA quality was assessed by A260/A280 ratio using a Nanodrop (Thermo Fisher Scientific), and integrity was confirmed by 1.5% agarose gel electrophoresis. RNA concentrations were quantified using a Qubit 3.0 Fluorometer with the Qubit RNA Broad Range Assay Kit (Q10210; Life Technologies). Stranded RNA-seq libraries were prepared from 1 µg of total RNA using the Fast RNA-Seq Lib Prep Module Kit (RK20306; ABclonal), and 200–500 bp PCR products were enriched, quantified, and sequenced on an Illumina Novaseq 6000 platform with 150 bp paired-end reads. Raw reads were filtered with fastp (v0.23.1), and clean reads were aligned to the human genome (GRCm38) using STAR with parameters “-sjdbScore 1 -outFilterMultimapNmax 20 -outFilterMismatchNmax 999 -outFilterMismatchNoverReadLmax 0.04 -alignIntronMin 20 -alignIntronMax

1000000 -alignMatesGapMax 1000000 -alignSJoverhangMin 8 -alignSJDBoverhangMin 1.” Gene-level counts were obtained using featureCounts (Subread v1.5.3) with the “-M” option, and differential expression analysis was performed with DESeq2. Genes with Fold change >2 and *P* < 0.05 were considered significantly differentially expressed. Heatmaps were generated from the intersection of differentially expressed genes across samples, and Gene Set Enrichment Analysis (GSEA) was conducted using the ClusterProfiler package (v3.14.3).

### RIP-seq

For RNA immunoprecipitation, Caco-2-N cells from RDPs-infected and mock-treated groups were harvested and lysed in 5 mL NP-40 buffer on ice. Clarified lysates were incubated with 45 µL IgG or PABPC1 antibody for 2 h at 4 °C. Protein A/G magnetic beads (80 µL per sample) were pre-washed three times with DEPC-treated PBST and added to the antibody–lysate mixtures, followed by overnight incubation at 4 °C with gentle rotation. Beads were washed four times with PBST and resuspended in 200 µL PBST. RNA was extracted by adding 800 µL TRIzol LS directly to the bead suspension. Input and immunoprecipitated RNA were purified and used for stranded RNA-seq library preparation.

### Nanopore poly(A)-tail sequencing

Poly(A)-tail length was measured using Oxford Nanopore direct RNA sequencing. Total RNA was extracted with TRIzol and treated with DNase I, and RNA integrity was confirmed before library preparation. Direct RNA libraries were prepared using the ONT Direct RNA Sequencing Kit following the manufacturer’s protocol and sequenced on R9.4.1 flow cells. Raw FAST5 files were basecalled with Guppy (high-accuracy mode), and reads were aligned to the reference genome using minimap2. Poly(A)-tail lengths were quantified using nanopolish polya or tailfindr, and only high-confidence tail-length calls were retained for downstream analyses. Poly(A)-tail distributions were calculated for each sample to compare host and viral RNAs

### Immunoprecipitation and mass spectrometry (IP–MS)

Caco-2-N cells were infected with RDPs and harvested at ∼36 h post-infection when robust GFP signals appeared. Cells from six wells were pooled per condition and lysed in NP-40 buffer on ice for 30 min, followed by clarification (12,000 rpm, 4 °C). The supernatant was divided equally and incubated for 2 h at 4 °C with gentle rotation with either 45 μg normal IgG or 45 μg target-specific antibody. Protein A/G magnetic beads (100 μL per sample), pre-washed with PBST and equilibrated in lysis buffer, were added to each mixture and rotated overnight (>8 h) at 4 °C. Beads were washed five times with PBST and three times with PBS, and bound complexes were subjected to on-bead reduction, alkylation and trypsin digestion. Resulting peptides were desalted using C18 StageTips and analysed by LC–MS/MS to identify proteins specifically enriched relative to IgG controls.

### Histopathological analysis

Histopathological analysis of mouse tissues was performed by Wuhan Servicebio Technology Co., Ltd. For histopathology, tissues were immediately fixed in 4% paraformaldehyde (PFA), embedded in paraffin, sectioned, and stained with hematoxylin and eosin (H&E).

### Statistical analysis

All data are presented as mean with SD, generated by GraphPad Prism 8.4.0. The statistical significance analyses were performed using a two-sided unpaired t test (P values) between two groups. *\*P* < 0.05, *\*\*P* < 0.01, *\*\*\*P* < 0.001, and *\*\*\*\*P* < 0.0001. P < 0.05 was considered statistically significant. Data distribution was assumed to be normal but this was not formally tested.

## Funding

This study was supported by grants from the National Key R&D Program of China (2021YFA1300800), China NSFC projects (82341061 and 82502690), and the Fundamental Research Funds for the Central Universities (2042022dx0003).

## Author contributions

Y.C., H.W., L.Z. and S.Y. conceived the research and experiments. S.Y., H.W., W.Y., S.Z., X.H., X.G. and J.L. performed the major experiments and data analysis. J.D. assisted with SARS-CoV-2 ΔN-GFP-HiBiT RDPs experiments. M.G., W. X., X.W., Z.Z. and Z.H. participated in cell and mice experiments of live SARS-CoV-2. K.L. and L.Z. provided critical advice. S.Y., H.W. and Y.C. wrote and revised the manuscript with contributions from all other authors.

## Declaration of interests

The authors declare no competing interests.

## Data and Materials Availability

RNA-seq, RIP-seq and Tail-seq data have been deposited in the Genome Sequence Archive (GSA) database under the accession number: HRA004999, HRA006231, HRA015381, CRA035193. RNA-seq datasets from other studies used in this work have been deposited in the NCBI Gene Expression Omnibus (GEO) under accession numbers GSE217504 and CRA002390. All other data are included in the article and supplemental information.

## Supplementary Materials

This PDF file includes: fig. S1 to S9, tables S1 and S2.

**Supplementary fig. S1.**
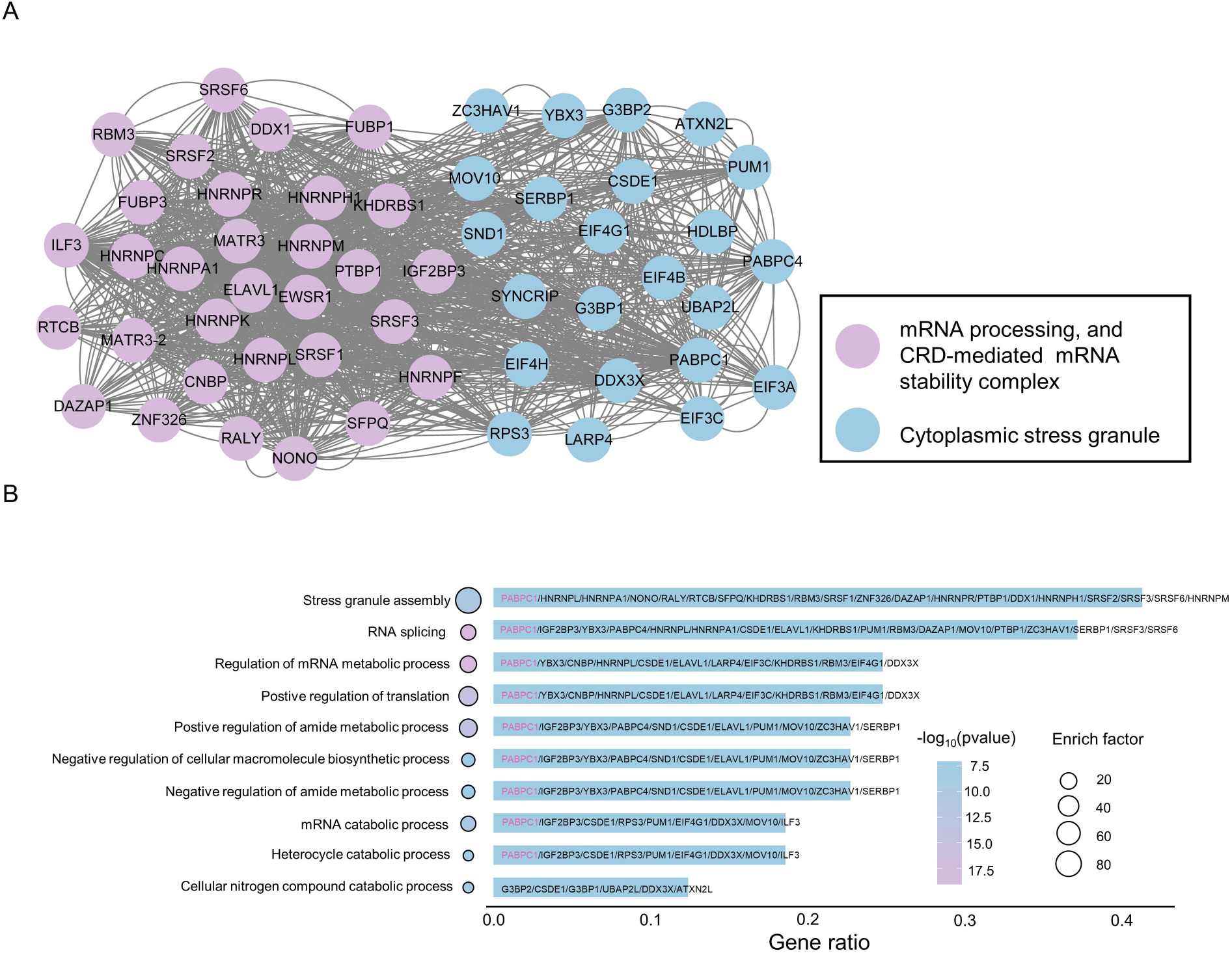
Functional clustering of high-confidence SARS-CoV-2 RBPs. (A-B) Protein-protein interaction (PPI) network analysis and GO enrichment of the 49 RBPs reveals clustering into mRNA metabolism and cytoplasmic stress granules.

**Supplementary fig. S2.**
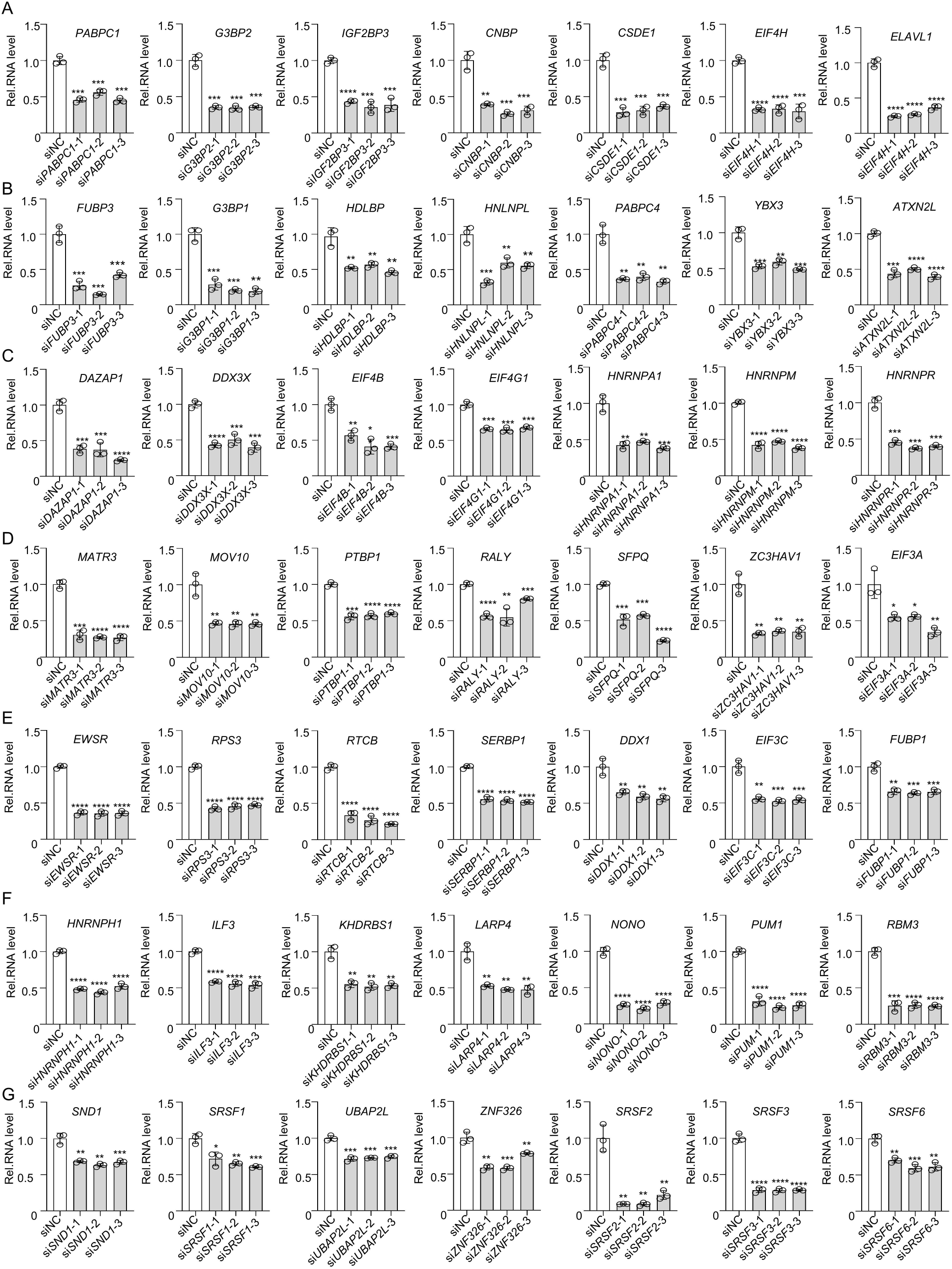
Validation of siRNA-mediated knockdown of RBPs. qPCR analysis confirming knockdown efficiency of three independent siRNAs targeting each of the 49 RBPs in Caco-2-N cells. Data are representative of three independent experiments and were analyzed by two-tailed unpaired t test. Graphs show the mean ± SD (n = 3) derived from three independent experiments. NS, not significant for *P* > 0.05, *\*P* < 0.05, *\*\*P* < 0.01, *\*\*\*P* < 0.001.

**Supplementary fig. S3.**
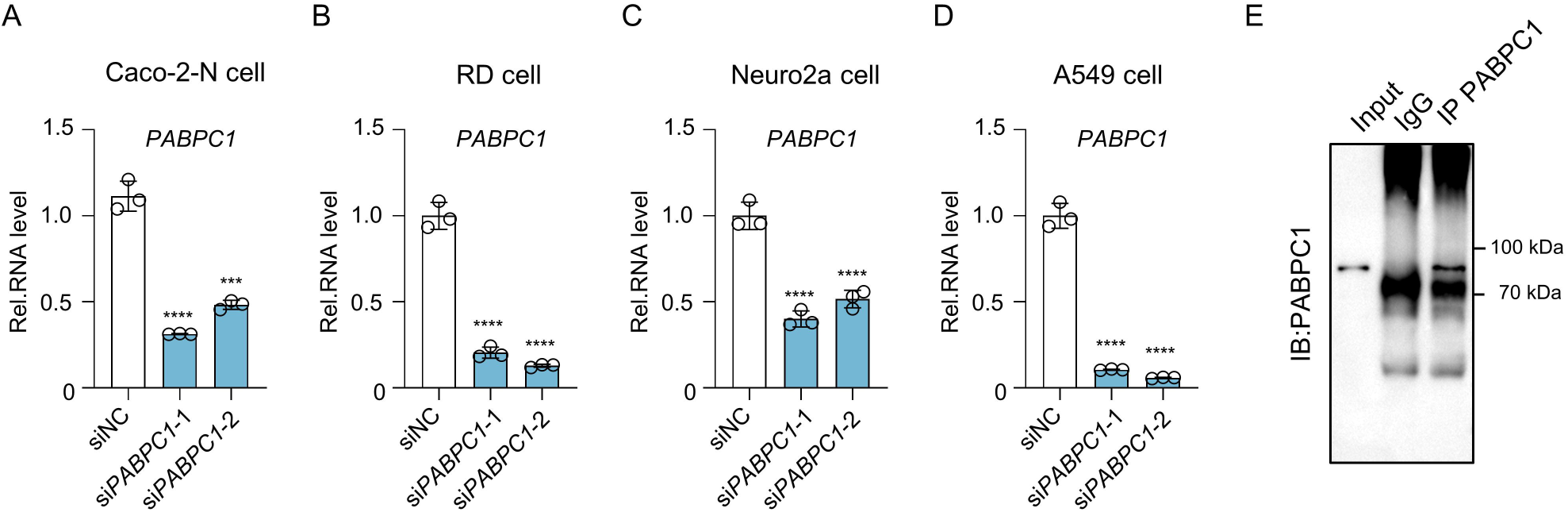
PABPC1 knockdown across several cell types. (A-D) qPCR analysis confirming PABPC1 knockdown in Caco-2-N, RD, Neuro2a and A549 cells. (E) Western blot analysis of input, IgG control, and PABPC1 immunoprecipitated (IP) samples. Data are representative of three independent experiments and were analyzed by two-tailed unpaired t test. Graphs show the mean ± SD (n = 3) derived from three independent experiments. NS, not significant for *P* > 0.05, *\*P* < 0.05, *\*\*P* < 0.01, *\*\*\*P* < 0.001.

**Supplementary fig. S4.**
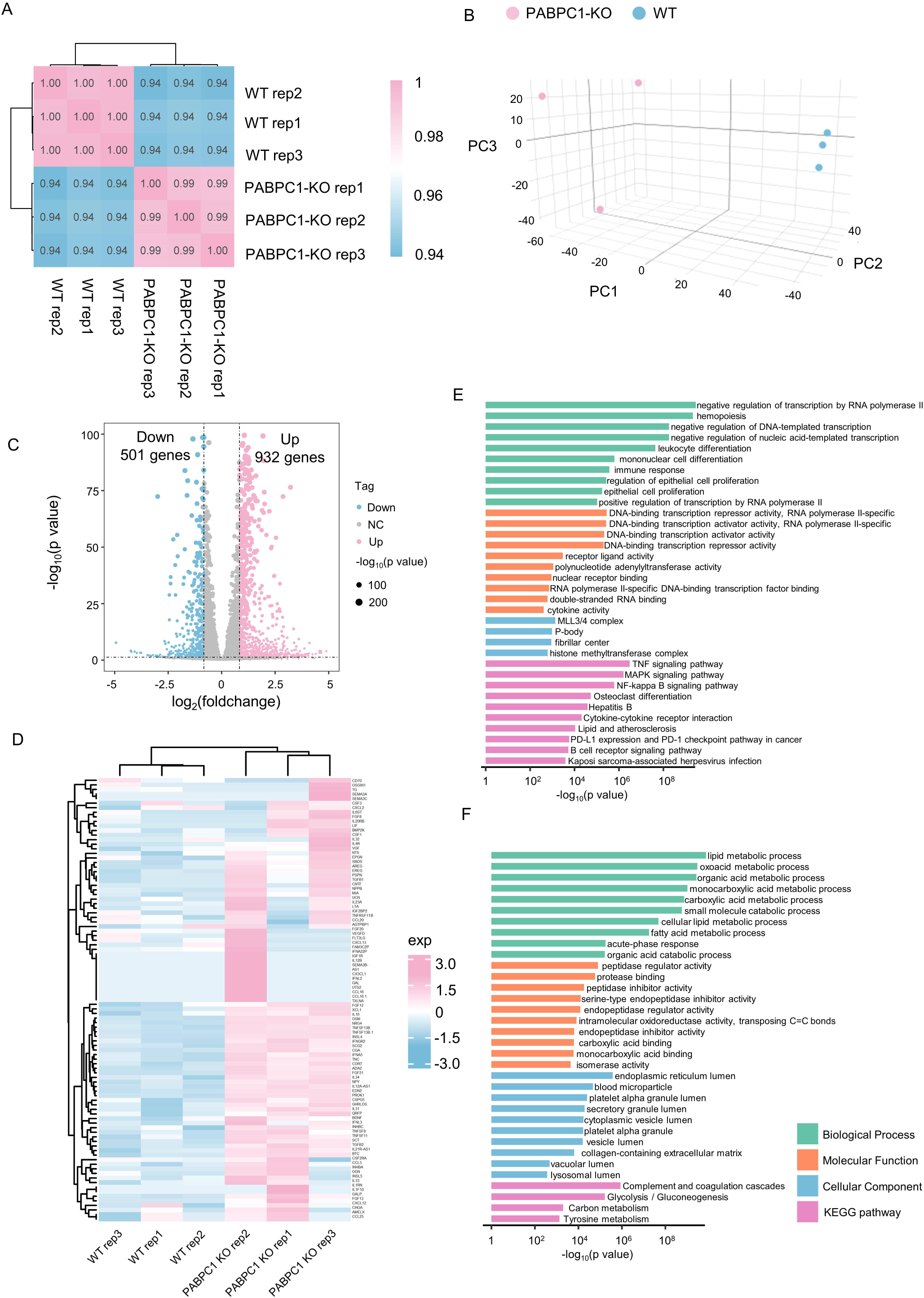
Transcriptomic impact of PABPC1 knockout. (A-B) Pearson correlation heatmap and principal component analysis (PCA) of gene expression in wild-type and PABPC1 knockout Caco-2 cells following SARS-CoV-2 infection. (C) Volcano plot analysis of differential gene expression in wild-type and PABPC1 knockout Caco-2 cells following SARS-CoV-2 infection. (fold change > 2, P < 0.05). (D) Heatmap of inflammatory cytokine expression in wild-type and PABPC1 knockout Caco-2 cells following SARS-CoV-2 infection. (E-F) GO and KEGG pathway enrichment analysis of upregulated and downregulated genes in wild-type and PABPC1 knockout Caco-2 cells following SARS-CoV-2 infection.

**Supplementary fig. S5.**
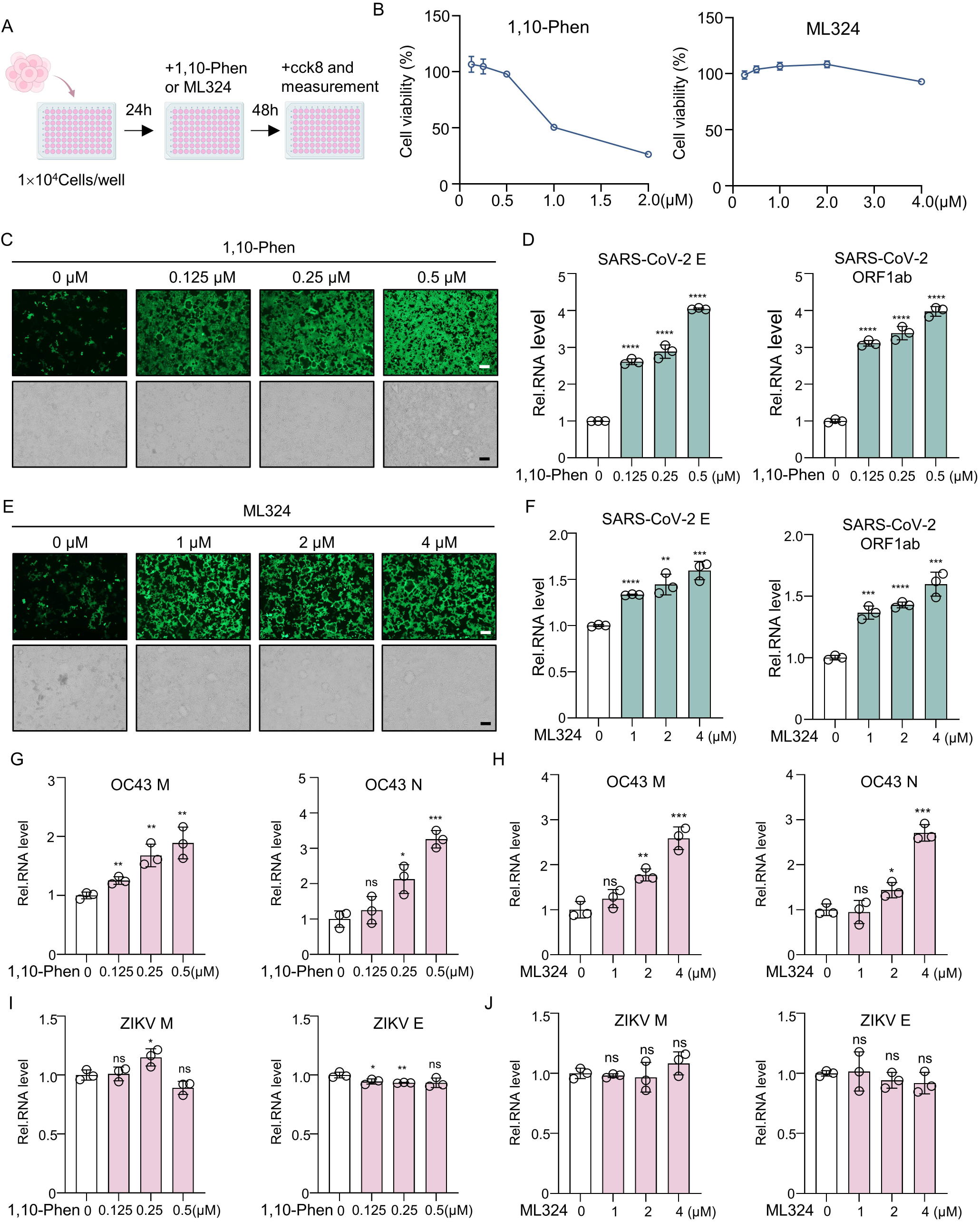
Effect of 1,10-Phen and ML324 on SARS-CoV-2 and other viruses. (A-B) Schematic representation of the CCK-8 assay showing minimal cytotoxicity of the small-molecule inhibitors 1,10-Phen and ML324. (C-D) GFP fluorescence and qPCR analysis of SARS-CoV-2 E and ORF1ab RNA levels in Caco-2-N cells pretreated with 1,10-Phen prior to infection with SARS-CoV-2 RDPs. (E-F) GFP fluorescence and qPCR analysis of SARS-CoV-2 E and ORF1ab RNA levels in Caco-2-N cells pretreated with ML324 prior to infection with SARS-CoV-2 RDPs. (G-H) qPCR analysis of HCoV-OC43 M and N RNA levels in Neruo2a cells pretreated with 1,10-Phen and ML324 prior to infection with HCoV-OC43. (I-J) qPCR analysis of ZIKV M and E RNA levels in Neruo2a cells pretreated with 1,10-Phen and ML324 prior to infection with HCoV-OC43. Data are representative of three independent experiments and were analyzed by two-tailed unpaired t test. Graphs show the mean ± SD (n = 3) derived from three independent experiments. NS, not significant for *P* > 0.05, *\*P* < 0.05, *\*\*P* < 0.01, *\*\*\*P* < 0.001.

**Supplementary fig. S6.**
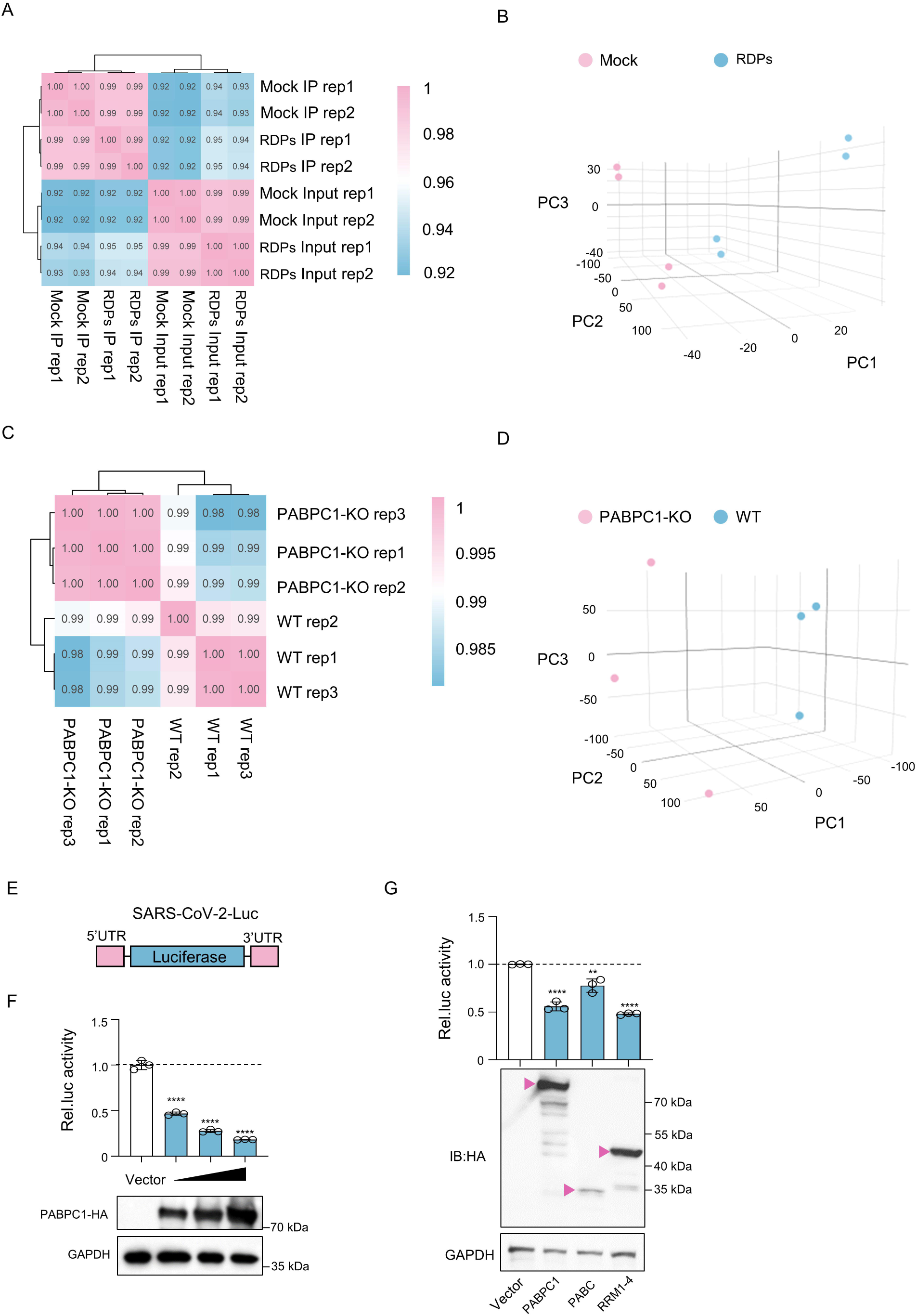
RIP-seq and poly(A) tail analysis of PABPC1. (A-B) Pearson correlation heatmap and principal component analysis (PCA) of gene expression in input and PABPC1 immunoprecipitated (IP) samples from Caco-2-N cells, with or without SARS-CoV-2 RDPs infection. (C-D) Pearson correlation heatmap and principal component analysis (PCA) of gene expression in wild-type and PABPC1 knockout Caco-2 cells following SARS-CoV-2 infection. (E) Schematic representation of SARS-CoV-2 Luciferase reporter. (F) Relative luciferase activity of the SARS-CoV-2-Luc reporter in cells transfected with 100 ng, 200 ng, or 500 ng of PABPC1-HA. (G) Relative luciferase activity of the SARS-CoV-2-Luc reporter in cells transfected with 500 ng of PABPC1 truncation fragments. Data are representative of three independent experiments and were analyzed by two-tailed unpaired t test. Graphs show the mean ± SD (n = 3) derived from three independent experiments. NS, not significant for *P* > 0.05, *\*P* < 0.05, *\*\*P* < 0.01, *\*\*\*P* < 0.001.

**Supplementary fig. S7.**
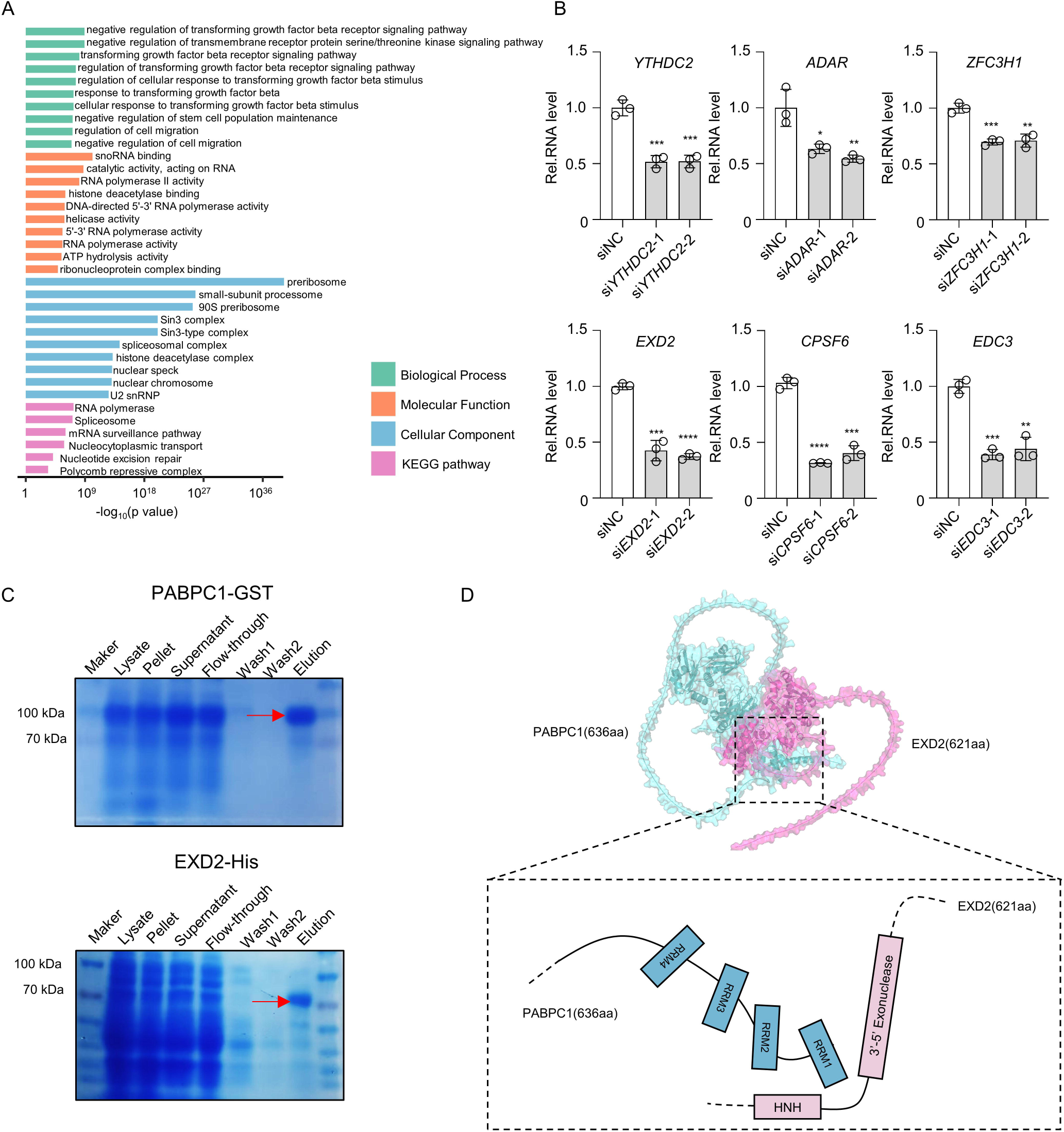
Identification of PABPC1-interacting proteins and structural mapping. (A) GO and KEGG enrichment of PABPC1 interactors in SARS-CoV-2 RDPs-infected Caco-2-N cells. (B) Validation of siRNA knockdown efficiency for candidate interactors. (C) Coomassie-stained SDS-PAGE showing purified PABPC1-GST and EXD2-His proteins. (D) AlphaFold3 prediction of PABPC1-EXD2 structural interface. Data are representative of three independent experiments and were analyzed by two-tailed unpaired t test. Graphs show the mean ± SD (n = 3) derived from three independent experiments. NS, not significant for *P* > 0.05, *\*P* < 0.05, *\*\*P* < 0.01, *\*\*\*P* < 0.001.

**Supplementary fig. S8.**
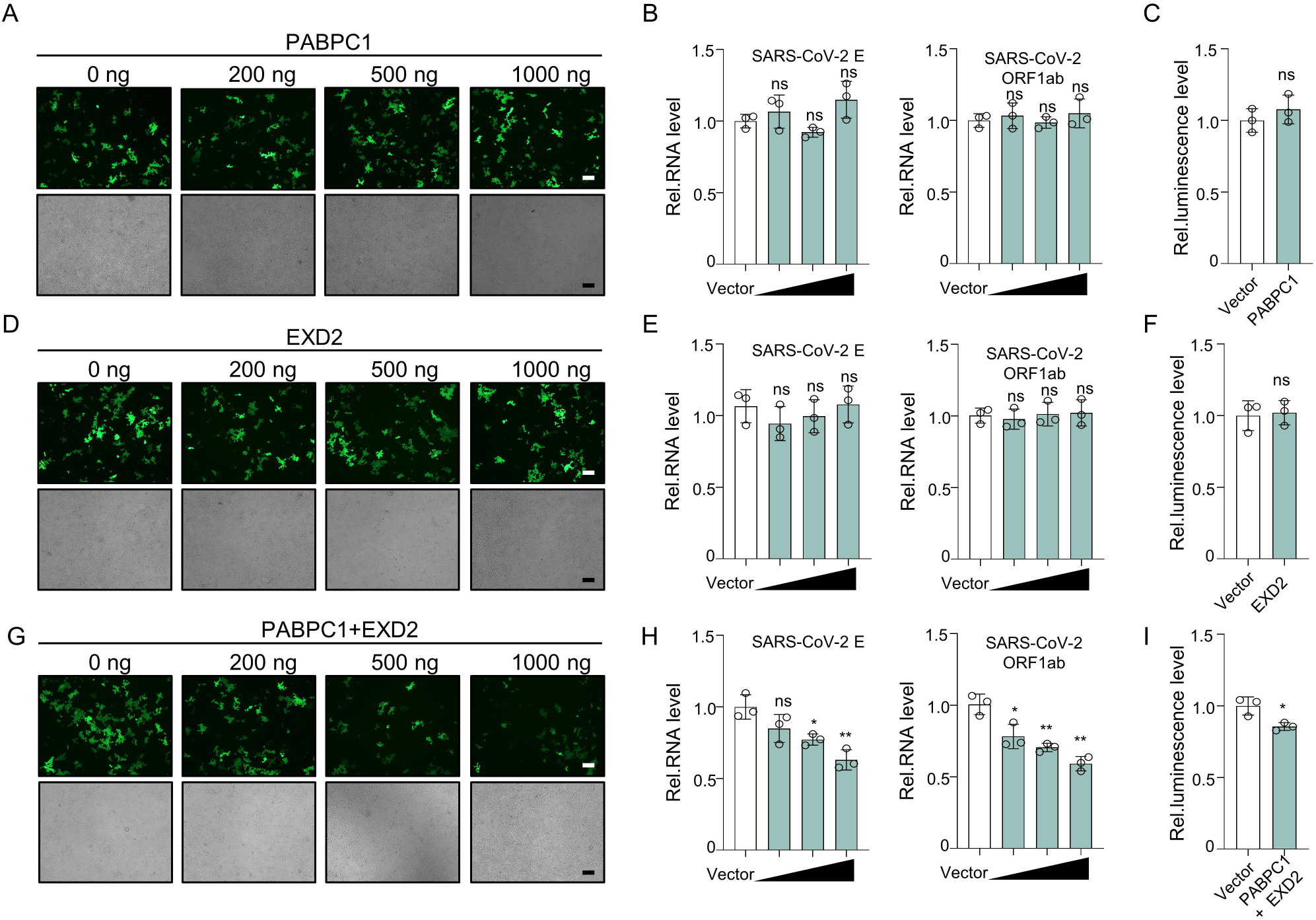
Co-expression PABPC1 and EXD2 domains inhibit viral replication. (A-C) qPCR analysis of SARS-CoV-2 E and ORF1ab RNA levels, GFP fluorescence, and kinetics of HiBiT luminescence in Caco-2-N cells following transfection with PABPC1 and infection with SARS-CoV-2 RDPs. (D-F) qPCR analysis of SARS-CoV-2 E and ORF1ab RNA levels, GFP fluorescence, and kinetics of HiBiT luminescence in Caco-2-N cells following transfection with EXD2 and infection with SARS-CoV-2RDPs. (G-I) qPCR analysis of SARS-CoV-2 E and ORF1ab RNA levels, GFP fluorescence, and kinetics of HiBiT luminescence in Caco-2-N cells following transfection with PABPC1 and EXD2 and infection with SARS-CoV-2RDPs. Data are representative of three independent experiments and were analyzed by two-tailed unpaired t test. Graphs show the mean ± SD (n = 3) derived from three independent experiments. NS, not significant for *P* > 0.05, *\*P* < 0.05, *\*\*P* < 0.01, *\*\*\*P* < 0.001.

**Supplementary fig. S9.**
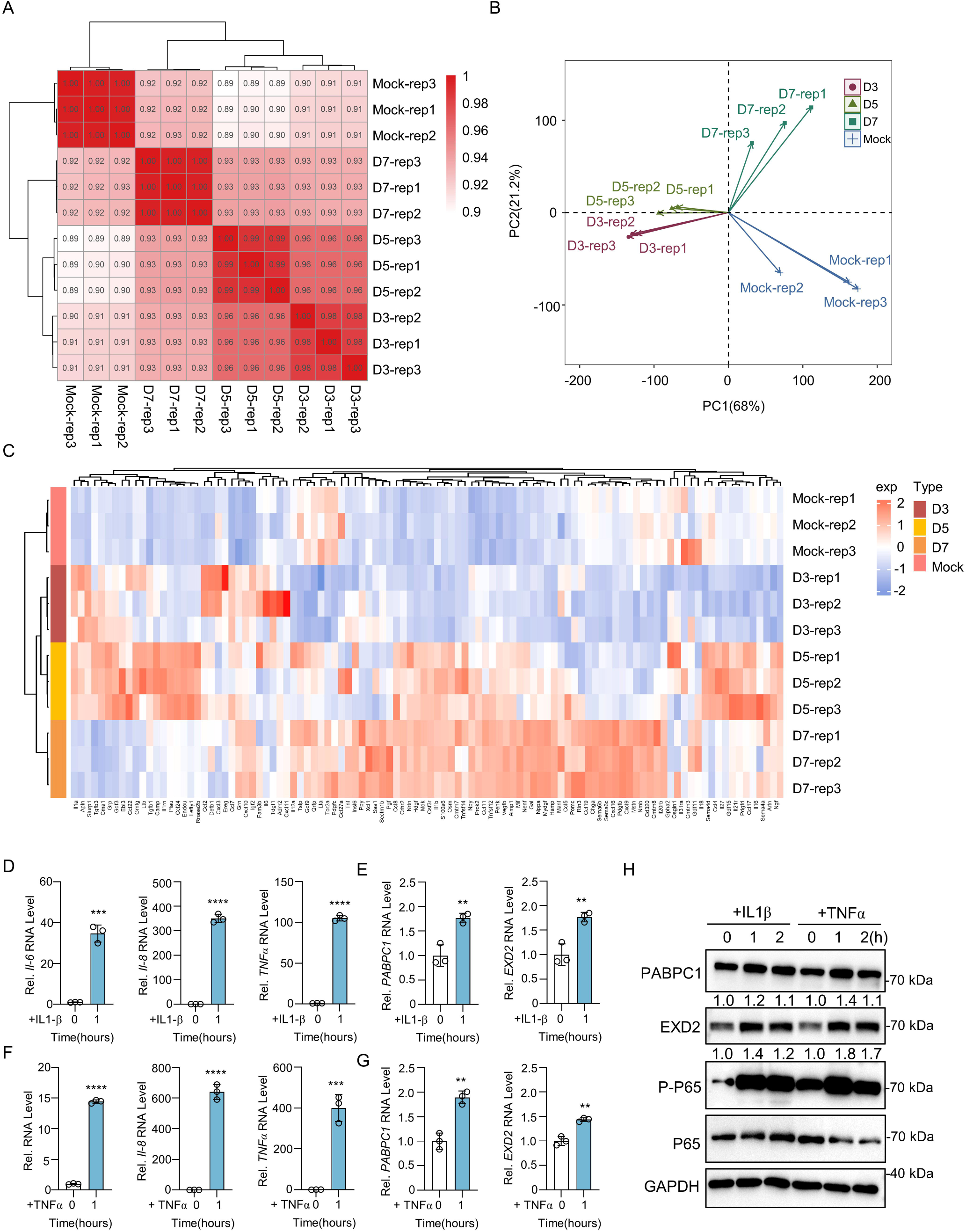
PABPC1 and EXD2 are upregulated during inflammatory response. (A-B) Pearson correlation heatmap and principal component analysis (PCA) of gene expression in the lungs of SARS-CoV-2-infected K18-hACE2 mice. (C) Heatmap of inflammatory cytokine expression in the lungs of SARS-CoV-2-infected K18-hACE2 mice at days 3, 5, and 7 post-infection. (D) qPCR analysis of IL6, IL8, and TNFα mRNA levels in A549 cells following treatment with IL-1β. (E) qPCR analysis of PABPC1 and EXD2 mRNA levels in A549 cells following treatment with IL-1β. (F) qPCR analysis of IL6, IL8, and TNFα mRNA levels in A549 cells following treatment with TNFα. (G) qPCR analysis of PABPC1 and EXD2 mRNA levels in A549 cells following treatment with TNFα. (H) Western blot analysis of PABPC1 and EXD2 protein levels in A549 cells following treatment with IL-1β or TNFα. Data are representative of three independent experiments and were analyzed by two-tailed unpaired t test. Graphs show the mean ± SD (n = 3) derived from three independent experiments. NS, not significant for *P* > 0.05, *\*P* < 0.05, *\*\*P* < 0.01, *\*\*\*P* < 0.001.

**Table S1.** siRNA sequences.

**Table S2.** Proteins identified as PABPC1 interactors by IP-MS.

## Notes

### Competing Interest Statement

The authors have declared no competing interest.

